# Metallothionein loss in cancer cells contributes to increased mutations through defective DNA repair and metabolic imbalance

**DOI:** 10.64898/2026.07.06.736843

**Authors:** Mirna Mina-Abouda, Amy C Rees, Della Evans, Evan Villamor, George Fullbright, Heather R Ghent, Madison A Clark, Wendy Y Zhang, Isabel Koehler, Isabella Berry, Sydney Oesch, Robert Hutchinson, Davide Delisi, Christopher de Solis, Alexander Y Maslov, Christopher Bradley, Samim Sharifi, Rosemarie Elloisa P Acero, Yuri K Peterson, Jie Zhang, Zhiwei Ye, Tori C Rodrick, Danyelle M Townsend, Saverio Gentile, Brian Orr, Drew Jones, Jessica H Hartman, David T Long, Jonathan T Sczepanski, Joe R Delaney

## Abstract

Understanding which genes are involved in mutagenesis is essential for developing cancer prevention and treatment strategies; establishing protectors of the genome has revolutionized cancer biology. Here, we describe metallothionein (MT) proteins as previously uncharacterized protectors against mutagenesis. MT is a heavy metal binding protein essential for zinc homeostasis and protection against heavy metal cytotoxicity. Because zinc binds approximately 10-15% of the proteome and is critical for processes such as DNA repair and mitochondrial health, MT loss is expected to disrupt these processes. We hypothesized that MT loss induces genomic instability by impairing DNA repair and mitochondrial function. In this study, the consequences of MT deficiency in high-grade serous ovarian cancer (HGSC) were investigated by knockdown of the most highly expressed MT, *MT2A*. Loss of *MT2A* resulted in the impaired DNA repair pathway base excision repair (BER), leading to increased mutagenesis. *MT2A* deficiency produced mitochondrial dysfunction, characterized by a decrease in mitochondrial membrane potential, glycolysis, oxidative phosphorylation, amino acids, and an imbalance of nucleobases. Together, these defects reflect cellular states associated with increased cancer aggressiveness. These findings identify MT as a fundamental hub maintaining genomic and metabolic integrity.

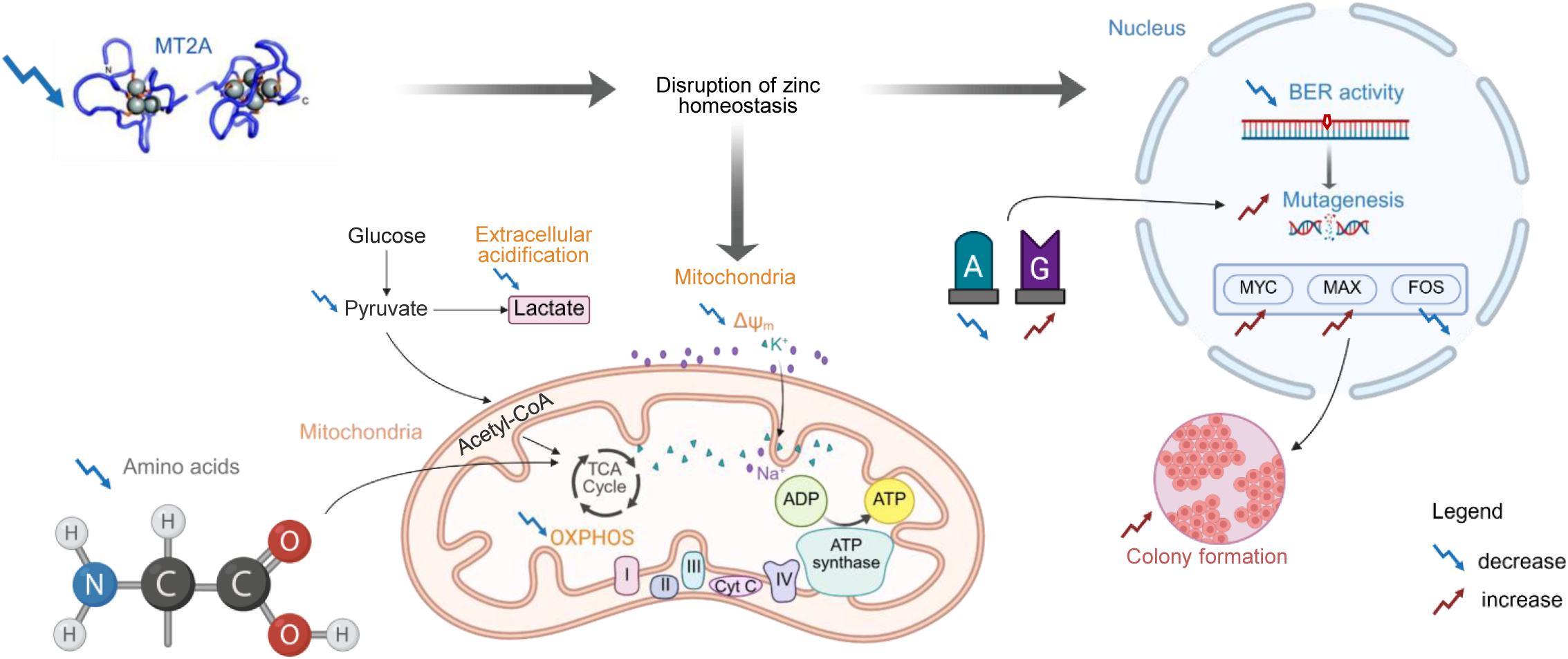

## Introduction

Ovarian cancer’s somatic mutation origins remain poorly understood. Ovarian cancer (OC) is a leading cause of gynecologic cancer deaths(1–3). OC is classified into three major types: epithelial, germ cell, and sex cord-stromal tumors. Epithelial OC, which accounts for most cases, includes four histologic subtypes: serous, endometrioid, mucinous, and clear cell carcinomas. Among these, serous carcinomas are further divided into low-grade (10%) and high-grade (90%) forms(2,4). High-grade serous ovarian cancer (HGSC) is an aggressive and lethal subtype, with a 5-year survival rate of only 29%(5). HGSC is characterized by mutations in the p53 (*TP53)* gene(6,7) and widespread aneuploidy(8–10), defined as the gain or loss of chromosomes or chromosome arms. While the contribution of *TP53* mutations to HGSC cancer initiation and progression has been extensively studied, the contribution of aneuploidy and other forms of copy-number alterations (CNAs) to cancer initiation and progression remains an emerging area of research.

Metallothioneins (MTs) are a gene cluster commonly affected by chromosome loss in cancer. We recently published data showing that 69% of HGSC cases lose the chromosome 16q13 genomic region encoding metallothionein (MT) proteins(11). MTs are a family of small (∼6-7 kDa), cysteine-rich, metal-chelating proteins with four major isoforms: MT1, MT2, MT3, and MT4. These isoforms are encoded by 11 adjacent genes: *MT4*, *MT3*, *MT2A*, *MT1E*, *MT1M*, *MT1A*, *MT1B*, *MT1F*, *MT1G*, *MT1H*, and *MT1X*(12,13). *MT2A* is the isoform most abundantly expressed in adult and cancer tissues with exceptions in skin (*MT3*) and brain (*MT4*)(14,15), suggesting MT2A may play a dominant role in regulating MT-associated functions. MTs serve as a major reservoir for labile zinc, a heavy metal that interacts with 10-15% of the proteome(16–18). Zinc-dependent proteins play critical roles in base excision repair (BER)(19) and homology-directed repair (HR)(20,21), which are DNA repair pathways responsible for repairing oxidative DNA lesions and DNA double-strand breaks, respectively. Additionally, zinc-dependent proteins share responsibility in maintaining mitochondrial integrity and bioenergetic function(22–25). Disruption of zinc homeostasis through MT gene loss has the potential to impair both nuclear genome maintenance and mitochondrial metabolism.

MTs bind toxic heavy metals such as cadmium and serve as a main protection mechanism against heavy metals. Cadmium exposure induces MT expression and displaces zinc from MT proteins, leading to altered zinc availability and conformational changes in MTs and other zinc-bound proteins(26–29). Cadmium is an established environmental carcinogen(30,31) and has been shown to induce DNA damage(17). Our previous work demonstrated that *MT2A*-deficient cells accumulate increased γH2AX foci following cadmium exposure(32). However, whether *MT2A* loss is sufficient to drive mutagenesis, with or without cadmium, remained unresolved.

Here, we establish for the first time MT2A as a mutagenesis protecting factor in human cells. We investigated the consequences of *MT2A* loss on genome stability, DNA repair fidelity, and mitochondrial function in HGSC. Using a controlled *MT2A* knockdown model that mimics the partial loss observed in patient tumors, we applied single-molecule mutation sequencing to directly measure random spontaneous mutations, independent of sequencing artifacts. Mutations can occur due to an increase in damage or a decrease in repair, which we measured through lesion-specific assays. We assessed base excision repair capacity, transcriptional programs associated with stemness, and mitochondrial morphology, bioenergetics, and metabolism. Together, these approaches allowed us to test the central hypothesis that *MT2A* loss drives genomic instability by disrupting metabolic and DNA repair homeostasis in HGSC.

## Materials and methods

### Cell culture

CAOV3, OvTrpMyc F318LOVi2, and HEK293T cells were used in this study. All cell lines were maintained at 37°C in a humidified incubator with 5% CO_2_ and cultured in complete media, which is RPMI-1640 with L-glutamine (Corning MT10040CV) supplemented with 10% fetal bovine serum (FBS) (Gibco A5256701), 1% sodium pyruvate (Millipore Sigma S8636), and 1% penicillin/streptomycin (Gibco 15070063). OvTrpMyc F318LOVi2 cells were originally derived from the left ovary of the C57BL/6-Trp53^em1(Ovgp1-Trp53*R270H-Myc)Jdel/^J(33) (OvTrpMyc)(Jax 040400) female mouse 318, then collected from a peritoneal tumor following the second intraperitoneal inoculation in wild-type C57BL/6J mice. The OvTrpMyc mouse model is constitutively driven by the *Ovgp1* promoter in the fallopian tube to express a dominant gain of function R270H mutation in *Trp53* and murine *c-Myc*.

### Lentiviral transductions and transfections

Stable knockdown of human *MT2A* or murine *Mt2* was achieved using shRNA-mediated lentiviral transduction. Lentiviral particles were produced in HEK293T cells by transfection with shRNA constructs targeting *MT2A* (MT2A-KD1 [target GCAAAGAGTGCAAATGCACTT], MT2A-KD2 [CTGCAAATGCAAAGAGTGCAA]), *Mt2* (Mt2-KD1 [GCAAACAATGCAAATGTACTT], Mt2-KD2 [GCAAATGTACTTCCTGCAAGA]), or a same-backbone non-targeting scrambled control (shScr, Addgene 1864), along with standard lentiviral packaging plasmids (Invitrogen K4975-00). Viral supernatant was collected 48-72 hours post-transfection. CAOV3 and OvTrpMyc F318LOVi2 target cells were exposed to 0.45µm polyethersulfone filtered (Sigma SLHPR33RS) viral supernatant in the presence of polybrene (Millipore Sigma TR1003G) to enhance transduction efficiency. Following a 6-hour incubation, the viral-containing media was replaced with fresh complete media. Stable cell populations were generated by antibiotic selection using puromycin (Gibco A1113803): 1µg/ml for CAOV3 and 2µg/ml for F318LOVi2. Six independent stable cell lines were established: CAOV3 shScr, MT2A-KD1, and MT2A-KD2, and OvTrpMyc F318LOVi2 shScr, Mt2-KD1, and Mt2-KD2. Cells were maintained in puromycin-containing media.

### Proliferation and cell count assays by nuclear count

For proliferation and cell count assays, including drug sensitivity measurements, nuclei counts were used for analysis. Drugs included cadmium chloride (CdCl_2_) (Millipore Sigma 202908), methyl methanesulfonate (Millipore Sigma 129925), and hydrogen peroxide (Millipore Sigma 216763). Cells were fixed and stained using Hoechst 33342 (Invitrogen H3570, working concentration of 0.23µg/mL) in 1% paraformaldehyde (PFA) (VWR 100504-858) in phosphate buffered saline (PBS). The cells were imaged using a BioTek Lionheart FX microscope and nuclei were quantified by user optimized and set metrics using Gen5 software. Data were exported and analyzed using Microsoft Excel.

### Reverse transcription quantitative PCR (RT-qPCR)

To extract RNA, the PureLink RNA Mini Kit (Invitrogen 12183018A) was used according to the manufacturer’s protocol. Briefly, lysis buffer containing 1% β-mercaptoethanol was added directly to cells. The cells were passed through a 26G needle (Fisher Scientific 14-826-15) in a 1mL syringe (Fisher Scientific 14-823-434) 4-6 times to homogenize. Subsequently, 70% ethanol was added and mixed thoroughly. The mixture was transferred to an RNeasy spin column placed in a collection tube and centrifuged at 12,000 x g for 30 seconds (this centrifugation speed and duration were used for all steps unless otherwise noted). Next, Wash Buffer 1 was added and centrifuged, followed by Wash Buffer 2 twice. The membrane was dried by centrifugation at 12,000 x g for 2 minutes. RNA was eluted by adding RNase-free water directly to the membrane and incubating at room temperature for 1-2 minutes. The column was then placed into a clean 1.5ml tube and centrifuged at 12,000 x g for 2 minutes to collect the RNA. RNA concentration was measured using a NanoDrop ND1000 spectrophotometer. RNA was stored at −20°C for short-term use and -80°C for long-term storage. To synthesize cDNA, the iScript cDNA synthesis kit (Bio-Rad 1708890) was used using the Bio-Rad protocol. Reverse transcriptase, iScript reaction mix, and RNA/water mixture were combined. Reverse transcription was performed at 25°C for 5 minutes, followed by 46°C for 20 minutes, and 95°C for 1 minute. cDNA was then stored at 4°C.To quantify the genes of interest, 400nM of primer (see Supplementary Table S1 for exact primer sequences) and 1X SYBR Green (Bio-Rad 172-5124), and cDNA were combined. Amplification was performed with an initial denaturation at 95°C for 30 seconds, followed by 39 cycles of 95°C for 5 seconds and 59°C for 30 seconds. Data were analyzed using the ΔΔCt method in Microsoft Excel.

### Immunofluorescence

Cells were fixed in 4% PFA in PBS for 10 minutes, followed by 0.1% Triton X-100 (Millipore Sigma X100) in PBS for 3 minutes. All incubations were at room temperature unless otherwise noted. The cells were blocked with 5% bovine serum albumin (BSA) (VWR 80055-682) and 5% goat serum (VWR 102643-594) in PBS for 45 minutes. MT2A primary antibody (Abcam ab192385) or gamma-H2AX (Biolegend 613401) were diluted 1:200 in blocking buffer overnight at 4°C. The cells were washed with PBS and anti-rabbit Alexa Fluor 488 secondary antibody (Invitrogen A11034) or anti-rat Alexa Fluor 647 secondary antibody (Invitrogen A212470) diluted 1:1000 in blocking buffer containing Hoechst 33342 (1:10,000) for 2 hours. After a series of PBS washes, ProLong Gold (Invitrogen P36930) was added, and the slides had a coverslip added and sealed by nail polish. The cells were imaged using the BioTek Lionheart FX microscope. Image analysis was performed using ImageJ and Microsoft Excel or Cellpose for punctae segmentation and then quantitation (34).

### SMM-sequencing and mutation spectrum analysis

CAOV3 cells transduced with either shScr or MT2A-KD were seeded into 6-well tissue culture treated plates. Each plate contained 2 wells of the same cell line: one was treated with 5µM CdCl_2_ and the other was not treated. All cells were cultured in complete RPMI. Once cells reached approximately 90% confluency, they were passaged by 1:10 cell dilutions. This was repeated for a total of 15 passages. After 15 passages, cells were transferred to a 10cm plate and harvested to isolate genomic DNA using the PureLink Genomic DNA Mini Kit (Invitrogen K182001). An additional sample was treated using 65µM of hydrogen peroxide for 3 passages to generate a control for DNA damage induced by reactive oxygen species (ROS). These cells were similarly moved to a 10cm plate after 3 passages and harvested to isolate gDNA. Genomic DNA samples were quantified using the Invitrogen Qubit 3.0 Fluorometer and the Qubitª dsDNA BR Assay Kit (Invitrogen Q32850) and then shipped to Mutagentech for library construction, sequencing, and analysis. Mutagentech then followed previously published methods for single-molecule mutation sequencing (SMM-seq), nonnegative matrix factorization mutation signature creation, and MutationalPatterns analysis(35). Results were aligned to hg19. All mutations are included as Supplementary Table S2, with a summary in Supplementary Table S3, and raw data have been deposited in SRA under Bioproject PRJNA1213734. Signature comparison to The Cancer Genome Atlas (TCGA) tumors utilized the file mc3.v0.2.8.PUBLIC.maf.gz in the Genomic Data Commons database as the input for MutationalPatterns(36). Novel signatures are supplied as Supplementary Table S4. Claude Code AI was used to develop and clean portions of this signature TCGA analysis workflow. A minimum of 50 mutations per sample was required for input into the analysis. Single-base substitution (SBS) reference patterns were from Alexandrov *et al.* 2020(37). Code associated with this study is available at https://github.com/jrdelaney/Published_Code/.

### Live cell base excision repair assessment

The protocol mirrored that of previous publications by the reagent inventors (38,39), with minor changes noted here. Cells were seeded into RPMI medium without antibiotics and maintained at 37°C in a humidified incubator with 5% CO_₂_. Chimeric BER probes were prepared in buffer containing sodium chloride (NaCl, VWR 97061-270), Tris (hydroxymethyl)aminomethane hydrochloride (Tris-HCl, Millipore Sigma Trizma base T1503 and hydrochloric acid 320331), Dl-Dithiothreitol (DTT, Millipore Sigma 43815), and Ethylenediaminetetraacetic acid disodium salt dihydrate (EDTA, Millipore Sigma E5134) (pH 8.0). Probes were annealed by heating and gradual cooling (38,39). Cells were transfected with BER detection probes (10 pmol) using RNAiMAX (Fisher Sci 13-778-075) in Opti-MEM (Thermofisher Sci 31985062). Transfection complexes were incubated for 10 minutes at room temperature and added dropwise to cells. Media was replaced with complete RPMI medium and imaging was performed starting at 4 hours (movie) or 24 hours (quantitation) post-transfection using the CY5 channel on a BioTek Lionheart LFX imaging system. Microsoft Excel was used for analysis and DelaneyApps Spectrum was used for data plotting.

### Zinc bound enzyme characterization

Supplementary Table S1 in Burger et al. was utilized to determine which proteins had bound zinc, as assessed by the N,N,N′,N′-Tetrakis(2-pyridylmethyl)ethylenediamine (TPEN) treatment group in HCT116 cells(40). Adjusted p-values were obtained from the source table for displayed statistics.

### Homology-directed repair assay in *Xenopus* extracts

DNA repair assays were performed using *Xenopus* egg extracts. Briefly, plasmid DNA was incubated in high-speed supernatant (HSS) at 15 ng/ml at 21° C for 20 minutes. Reactions were then mixed with two volumes of nucleoplasmic extract (NPE) at 21° C to promote replication of plasmid DNA. 20 minutes after initiating replication, reactions were supplemented with buffer or 0.785 mM TPEN (N,N,N′,N′-tetrakis(2-pyridinylmethyl)-1,2-ethanediamine) (Millipore Sigma P4413). 40 minutes after initiating replication, reactions were supplemented with EcoRI (ThermoFisher Scientific ER0271), which cleaves a single recognition site on the plasmid to create a DNA double-strand break (DSB). At the times indicated, reaction samples were withdrawn, treated with 29.2 μM RNase (ThermoFisher Scientific EN0531) for 60 minutes at 21° C, and then treated with 13.3 μM proteinase K (Millipore Sigma 3115879001) for 16 hours. Samples were resolved by 0.8% agarose gel electrophoresis at 150 V for 2 hours. Gels were stained with SYBR Gold (Invitrogen S11494) and visualized using the ChemiDoc imaging system (Bio-Rad).

### Mouse cancer experiments

J:NU homozygous nude mice (Jax 007850) and C57BL/6J wild-type mice (Jax 000664) were used. Nude mice were injected intraperitoneally with 10 million CAOV3 shScr, MT2A-KD1, or MT2A-KD2 cells per mouse. Cell identities were blinded to the observer prior to injections, and blinding was maintained until completion of the experiment. Mice were monitored twice weekly for changes in weight and physical condition. Humane endpoints were defined as ≥10% weight gain or loss, or signs of physical distress. Mice meeting endpoints were euthanized via CO_2_ inhalation for 4 minutes, followed by secondary cervical dislocation. Tumors were harvested, flash frozen with liquid nitrogen, stored in liquid nitrogen, and submitted for metabolomic analysis. Wild-type mice were used to generate the OvTrpMyc F318LOVi2 cells as described under the cell culture section. All animal experiments were approved by the Medical University of South Carolina Institutional Animal Care and Use Committee under protocol #01305.

### Transcriptomic analysis

CAOV3 cells with lentiviral, integrated shScr or MT2A-KD were grown in complete RPMI. At 80% confluency, adherent cells were harvested for RNA using Invitrogen PureLink RNA kits (Invitrogen 12183018A). The Translational Science Laboratory at the Medical University of South Carolina utilized manufacturer protocols for polyA-based transcriptome library preparation (NEBNext® Ultra™ II Directional RNA Library Prep with Sample Purification Beads (NEB#E7490), polyA mRNA workflow). Libraries were sequenced at the Vanderbilt Genomics core using an Illumina NovaSeq X sequencer. Raw FASTQ files were aligned to hg38 using STAR and transcripts were counted using featureCounts. Differential expression analysis was performed using edgeR using the Galaxy platform(41). For transcription factor analysis, ChEA3 was used, using ChIP-seq data from the ENCODE project(42). Raw sequencing data are available within SRA under Bioproject PRJNA1428857 and raw count data are deposited in GEO under series GSE322490.

### Soft agar colony formation assay

A 2% low melting point agarose (IBI Scientific IB70051) solution was prepared in deionized water and autoclaved. The bottom layer consisted of 0.5% agarose prepared by diluting the 2% agar in complete media. A total of 50μL of the 0.5% agar was added per well of a 96-well plate and incubated at 37°C overnight to allow solidification. The top layer consisted of 0.3% agar diluted in complete media. The agar was reheated to liquefy and cooled to 45°C in a water bath prior to use. Cells were seeded at a density of 100 cells per well in the 0.3% agar top layer. An additional 50μL of complete media was added on top of the 0.3% agar layer. Plates were wrapped in parafilm and incubated at 37°C with 5% CO_₂_. Media was replenished every 2-3 days for 5-14 days until colonies formed, with equal incubation times maintained across the compared groups. Colonies were fixed using 0.005% crystal violet (Millipore Sigma 61135) in methanol (Millipore Sigma 34860), imaged, and quantified. Image analysis was performed using ImageJ, and data were analyzed using Microsoft Excel. Spectrum Data Plotter(43) was used for graph plots.

### Confocal microscopy of mitochondrial morphology

Cells were seeded at a density of 10,000 cells per well in Ibidi 18-well slides (Ibidi 81817) and allowed to adhere overnight. Cells were incubated for 30 minutes at 37°C with 5% CO_₂_ in media containing 75 nM MitoTracker Red CMXRos (Invitrogen M7512) and Hoechst 33342 at a 1:10,000 dilution. The staining media was aspirated, cells were washed with PBS, and fixation was performed using 4% PFA in PBS for 10 minutes at room temperature in the dark. PFA was aspirated, wells were washed with PBS, and cells were stored in PBS prior to imaging. Imaging was performed using a Zeiss LSM880 NLO with Airscan microscope at the Medical University of South Carolina Cell and Molecular Imaging Core. Images were acquired at 63× magnification with 4× zoom. Analysis was performed using the FIJI MitoTracker Analyzer plugin. The 2D threshold function was applied using the weighted mean method with a 1.25μm block size.

### Flow cytometry

CAOV3 and OvTrpMyc F318LOVi2 cells were seeded overnight in 6-well plates. Cells were incubated with Hoechst 33342 at a 1:10,000 dilution and tetramethylrhodamine methyl ester (TMRM) (VWR 115532-50-8) for mitochondrial membrane potential or 50mM Bis-(1,3-Dibutylbarbituric Acid)Trimethine Oxonol DiBAC4(3) (Invitrogen B348) for plasma membrane polarization for 30 minutes at 37°C. Control samples were treated with 100 μM carbonyl cyanide m-chlorophenyl hydrazone (CCCP) (Abcam Ab141229), a mitochondrial uncoupler. Cells were trypsinized and centrifuged at 1,000 × g for 2 minutes. Cell pellets were resuspended in cold PBS and centrifuged again. Pellets were resuspended in cold PBS and immediately analyzed using a Fluorescence-Activated Cell Sorting (FACS) Aria III flow cytometer. The median fluorescence intensity was analyzed using FlowJo and Microsoft Excel.

### Resipher oxygen consumption rate

CAOV3 cells were seeded at 5,000 cells per well using the Nunclon Delta-surface 96-well flat-bottom plates (ThermoFisher Scientific 167008) overnight in 37°C, 5% CO_2_. Resipher (Lucid Scientific RS-NS96) technology was used to measure oxygen consumption rate in 37°C, 5% CO_2_ incubator for a minimum of 4 hours. PFA and Hoechst were added, imaged via Lionheart LFX, and cells were counted by Gen5 software for normalization to nuclei count. Analysis was conducted using Microsoft Excel and a t-test was utilized to obtain statistics.

### Seahorse metabolic assays

For extracellular acidification rate measurements, 200 μL per well of Seahorse XF96 Calibrant solution (Agilent 100840-000) was added to the cartridge and incubated overnight at 37°C in a non-CO_₂_ incubator. CAOV3 cells were seeded at 7,500 cells per well in a Seahorse 96-well plate (Agilent 103794-100) and incubated overnight at 37°C with 5% CO_₂_. Media was aspirated, and cells were washed with PBS. RPMI XF assay media (Agilent 103576-100) supplemented with 1× glucose, 1× glutamine, and 1× sodium pyruvate (Agilent 103577-100, 103579-100, 103578-100) was added. Cells were incubated for 2 hours at 37°C in a non-CO_₂_ incubator prior to analysis. Glucose (10 mM) was loaded into port A, oligomycin (1 μM) into port B, and 2-deoxyglucose (10 mM) into port C (Cayman chemicals 14325).. The plate was calibrated and run using WavePro software. Immediately following analysis, cells were fixed with PFA and stained with Hoechst for normalization by nuclear counting. Nuclei were imaged using the Agilent Lionheart FX system and quantified automatically using Gen5 software.

### Metabolomics

Tumors were rapidly extracted from mice and snap frozen in liquid nitrogen. 10-20mg samples were isolated on dry ice and kept at -80°C prior to analysis. Samples were then moved from -80°C storage to wet ice and thawed. Extraction buffer proportional to mg of tumor was added, consisting of 80% methanol (Fisher Scientific # A412SK-4) and 500 nM metabolomics amino acid mix standard (Cambridge Isotope Laboratories #MSK-A2-1.2), was prepared and placed on dry ice. Samples were extracted by tumor pellets with extraction buffer (10mg/mL) in 2.0 mL screw cap vials containing ∼100 µL of disruption beads (Research Products International #9830). Each sample was homogenized for 10 cycles on a bead blaster homogenizer (Benchmark Scientific #D2400). Cycling consisted of a 30 sec homogenization time at 6 m/s followed by a 30 sec pause. Samples were subsequently spun at 21,000 g for 3 min at 4°C. A set volume of each (225 µL) was transferred to a 1.5 mL tube and dried down by vacuum centrifugation (Thermo Fisher #SpeedVac). Samples were reconstituted in 25 µL of Optima liquid chromatography (LC) mass-spectrometry (MS) grade water (Fisher Scientific #W6500). Samples were sonicated for 2 mins, then spun at 21,000 g for 3 min at 4 °C. 20 µL were transferred to LC vials containing glass inserts for analysis. The remaining sample was placed in -80 °C for long term storage.

Samples were subjected to an LC/MS analysis to detect and quantify known peaks. A metabolite extraction was carried out on each sample based on a previously described method(44). The LC column was a Millipore™ ZIC-pHILIC (2.1 x150 mm, 5 μm) coupled to a Dionex Ultimate 3000™ system. The column oven temperature was set to 25°C for the gradient elution. A flow rate of 100 μL/min was used with the following buffers; A) 10 mM ammonium carbonate (Sigma #207861) in water, pH 9.0, and B) neat acetonitrile (Sigma PHR1551). The gradient profile was as follows; 80-20% B (0-30 min), 20-80% B (30-31 min), 80-80% B (31-42 min). Injection volume was set to 2 μL for all analyses (42 min total run time per injection). MS analyses were carried out by coupling the LC system to a Thermo Q Exactive HF™ mass spectrometer operating in heated electrospray ionization mode (HESI). Method duration was 30 min with a polarity switching data-dependent top-5 method for both positive and negative modes. Spray voltage for both positive and negative modes was 3.5kV and capillary temperature was set to 320°C with a sheath gas rate of 35, aux gas of 10, and max spray current of 100 μA. The full MS scan for both polarities utilized 120,000 resolution with an AGC target of 3E6 and a maximum IT of 100 ms, and the scan range was from 67-1000 m/z. Tandem MS spectra for both positive and negative mode used a resolution of 15,000, AGC target of 1E5, maximum IT of 50 ms, isolation window of 0.4 m/z, isolation offset of 0.1 m/z, fixed first mass of 50 m/z, and 3-way multiplexed normalized collision energies (nCE) of 10, 35, 80. The minimum AGC target was 1e4 with an intensity threshold of 2E5. All data were acquired in profile mode.

The resulting Thermo^TM^ RAW files were converted to SQLite format using an in-house python script to enable downstream peak detection and quantification. The available MS/MS spectra were first searched against the NIST17 MS/MS(45), METLIN(46) and respective Decoy spectral library databases using an in-house data analysis python script adapted from our previously described approach for metabolite identification false discovery rate control (FDR)(47,48). Metabolite peaks were extracted based on the theoretical *m*/*z* of the expected ion type, e.g., [M+H]^+^, with a 15 part-per-million (ppm) tolerance and a ± 0.2 min peak apex retention time tolerance within an initial retention time search window of ±0.5 min. Statistics and heat maps were performed in Microsoft Excel by comparing shScr samples to all MT2A-KD tumors by t-test.

### Western blot

Cells were lysed in lysis buffer consisting of 60 mM Tris-HCl (pH 6.8), 12% sodium dodecyl sulfate (SDS), 47% glycerol, 12.5% β-mercaptoethanol, and 0.03% bromophenol blue supplemented with 1× Halt protease inhibitor (ThermoFisher Scientific 78443) and incubated for 10 minutes at room temperature. Cells were scraped, transferred to 1.5 ml tubes, and centrifuged at 13,000 x g for 10 minutes at 4°C. The supernatant was collected, boiled for 5 minutes at 95°C, and cooled to room temperature. Protein concentration was determined using a BCA assay, and equal amounts of protein were loaded onto Bio-Rad pre-cast gradient gels (Bio-Rad 4561093) for electrophoresis in 1× Tris/Glycine/SDS running buffer (Bio-Rad 1610772) prepared in deionized water. Wet transfer was performed onto polyvinylidene fluoride (PVDF) membranes (Millipore Sigma IPFL00010) using 1× Tris/glycine transfer buffer (Bio-Rad 1610771) containing 20% methanol for 45 minutes at 80 volts. Membranes were washed in PBS for 5 minutes and then blocked in 5% milk in PBS for 30 minutes at room temperature while rocking. Membranes were rinsed twice in tris-buffered saline with 0.1% tween-20 (TBST) and incubated with primary antibody OGG1(Proteintech 15125-1-AP) or Histone H2B (Invitrogen MA524697) diluted 1:1,000 in TBST for 1 hour at room temperature with rocking. Membranes were washed twice in TBST for 5 minutes each, followed by incubation with HRP-conjugated secondary antibody (Millipore Sigma AP307P and 12-349) diluted 1:5,000 in TBST for 30 minutes at room temperature with rocking. Membranes were washed in TBST with two quick rinses, followed by three 5-minute washes and one 30-minute wash. Enhanced chemiluminescence reagents (Bio-Rad 1705060) were mixed at a 1:1 ratio, applied to the membrane, and the signal was detected using a ChemiDoc imaging system.

### 8-oxo-dG ELISA

Genomic DNA damage was quantified using the EpiQuik 8-OHdG DNA Damage Quantification Direct Kit (Colorimetric) (EpigenTek P-6003-48). 600 ng of purified DNA from each sample was loaded into a 96-well plate according to the manufacturer’s instructions. Following completion of the assay protocol, absorbance at 450 nm was measured to determine relative levels of 8-OHdG.

### Scratch wound assay

Cells were seeded at 100,000 cells per well in a 24-well plate and incubated overnight at 37°C, 5% CO_2_. Wells were scratched with a 10µL pipette tip imaged with a BioTek Lionheart FX microscope from 0-24 hours. The scratch wound area was quantified using ImageJ and analyzed through Microsoft Excel.

## Results

### Modeling metallothionein genetics of ovarian cancer

To understand the contributions of MTs in cancer mutagenesis, we chose HGSC as a model system due to its moderate levels of mutations(37) even in the absence of other well-known tumor suppressors(7). HGSCs lose their MT gene cluster by copy-number alterations (CNA) loss at a much greater frequency than gaining the cluster: 90% of CNAs are losses and 10% are gains (**Fig. 1A**). These MT gene cluster deletions may be conferring a more chemo-resistant or aggressive phenotype, as monoallelic loss tumors have poorer 5-year overall survival prognosis (**Fig. 1B**). According to the TCGA PanCancer Atlas, less than 1% of tumors are marked as homozygous MT deletions, which may be an overestimate from technical or analytical error. Monoallelic loss contributes to *MT2A* mRNA decrease in these patient tumors (**Fig. 1C**). To model the loss case, a panel of HGSC cell lines within the Cancer Cell Line Encyclopedia (CCLE) was queried for mRNA levels of *MT2A*. CAOV3 cells expressed the most *MT2A* (**Fig. 1D**). Our previous research indicated that CAOV3 cells are uniquely useful model cells for MT biology in that they do not express all 11 metallothionein genes, but rather only two of them: *MT2A* and *MT1X* (32). *MT2A* is expressed 10-fold higher than *MT1X* in CAOV3 in our lab and others’ (32,49) and is consistently the highest expressed MT. This feature makes CAOV3 cells uniquely amenable to genetic manipulation for controlled experimentation of MT loss. A knockdown method is appropriate to transform CAOV3 to mimic the ovarian cancers with gene loss. We created two shRNA knockdown lines with independent shRNAs (MT2A-KD1 and MT2A-KD2). We noted the expected functional effects of *MT2A-*knockdown (MT2A-KD): increased sensitivity to CdCl_2_ (**Fig. 1E**). Cadmium is a heavy metal normally chelated by MT, so cell loss is consistent with effective depletion of MT activity. Functional changes in cadmium sensitivity correlated with the amount of MT knockdown, which was stronger in MT2A-KD1 than MT2A-KD2 (**Fig. 1F**). The depletion did not change the localization of MT2A; MT2A protein remained distributed throughout both the cytoplasm and nucleus in MT2A-KD (**Fig. 1G**).

**Figure 1.**
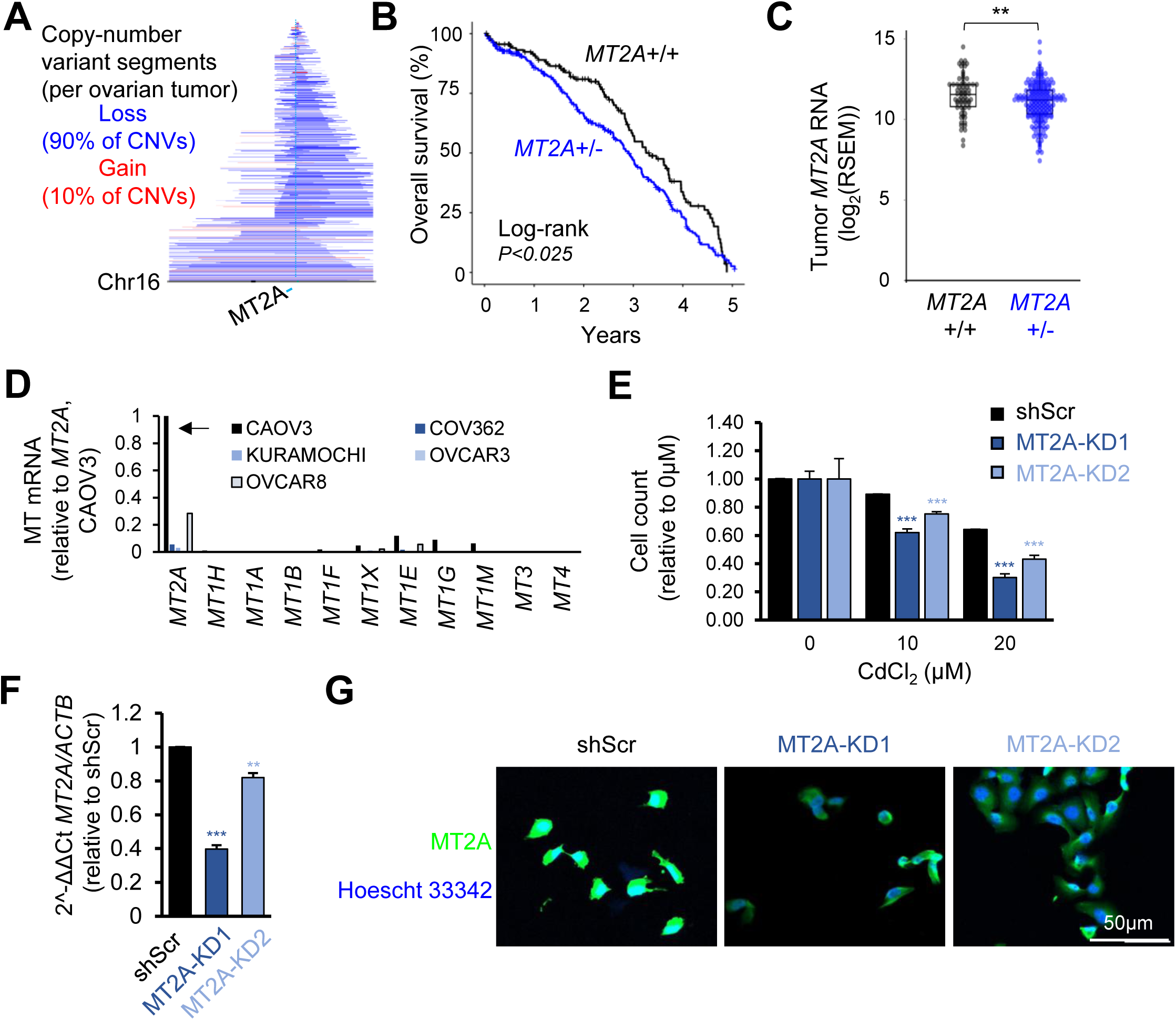
Modeling *MT2A*-deficiency in HGSC. (A) CAIRN plot of copy-number variants present in TCGA HGSC which overlap the MT gene cluster on chromosome 16. (B) TCGA PanCancer Atlas HGSC overall survival data of heterozygous *MT2A* loss tumors compared to tumors without *MT2A* loss. (C) Bulk RNA expression data from TCGA PanCancer Atlas HGSC tumors with normal *MT2A* copy number or without *MT2A* loss. (D) Bulk mRNA expression of all *MT* genes relative to CAOV3 *MT2A* levels in a panel of HGSC lines. (E) Phenotypic confirmation of *MT2A* knockdown through cellular proliferation by nuclei count (Hoechst 33342) assay following a 72-hour CdCl_2_ insult. (F) RNA level knockdown confirmation of *MT2A* by RT-qPCR. (G) Immunofluorescence of MT2A in control or knockdown cells to evaluate subcellular distribution. **P<0.01, ***P<0.001, by t-test.

### Ultra-low frequency mutation quantitation of low *MT2A* and cadmium treated HGSC with SMM-seq

Since a major known function of MTs is to sequester cadmium, one hypothesis supporting why cancers lose a copy of MT genes is that this enables environmental cadmium to further exert its mutagenic function. As an International Agency for Research on Cancer (IARC) regulated oncogenic agent, cadmium may promote tumorigenesis through mutational processes. Cadmium is a known carcinogen elevating the risk of lung cancer due to inhalation toxicity(30), but such toxic exposures are often confounded by other chemicals in smoke. We previously observed increased DNA damage signaling by gamma H2AX in MT knockdown models insulted with cadmium(32), but it remained unclear if damage went unrepaired and, if so, what spectrum of mutations might occur. To evaluate if mutations were produced by the interaction of low MT and the presence of low-dose cadmium, sublethal 5µM cadmium or vehicle control (water) was used to serially passage cells with shScr control or MT2A-KD constructs (**Supplementary Fig. S1A**). Genomic DNA was collected after 15 passages. A control for H_2_O_2_ induced mutations was conducted for 3-passages in parallel, at 65µM. To directly quantify random mutations within the genomic DNA, a cutting edge error correcting sequencing method, single-molecule mutation sequencing (SMM-seq), was used. SMM-seq utilizes DNA duplexes and rolling circle amplification methods to improve the detection sensitivity of ultra-rare spontaneous mutations by orders of magnitude compared to standard deep sequencing(35). This method is effective at subtracting out spurious mutations incurred by library creation, sequencer errors, or single-stranded damage, leaving bona-fide original genomic DNA mutations resulting from random mutagenesis (**Fig. 2A**). Overall, spontaneous single-nucleotide variants (SNVs) were increased in MT2A-KD cells relative to shScr (**Fig. 2B**), with an increase in CdCl_2_-moderated mutation only observed in the control shScr condition (**Supplementary Fig. S1B**). Thus, MT2A-KD dominated the observed mutational burden, rather than cadmium. This was counter to the original hypothesis that MT loss occurs in tumors to sensitize them to cadmium-mediated environmental mutagenesis, rather indicating MT biology itself is driving a tumor suppressor mutagenic phenotype with MT loss.

**Figure 2.**
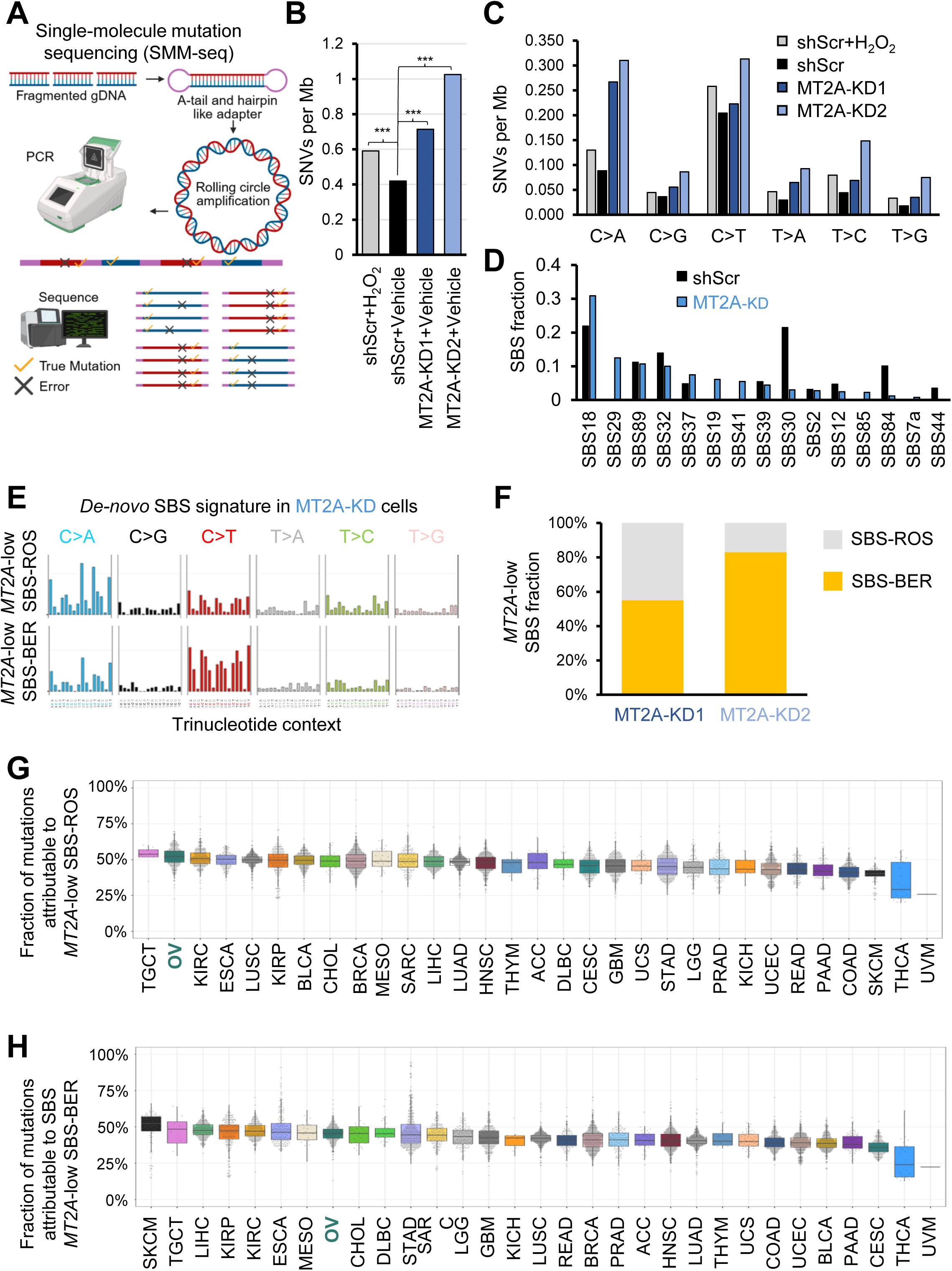
Metallothionein reduction incurs spontaneous mutagenesis. (A) Diagram of SMM-seq workflow and analysis, starting from genomic DNA (gDNA). (B) Total SNVs per megabase (Mb) of DNA analyzed by SMM-seq from the indicated CAOV3 cell groups. \*\*\**P*<0.001 by Fisher’s exact test. (C) Specific nucleotide changes SNVs per Mb of DNA analyzed by SMM-seq. Note that due to conventional reporting, C>A is equivalent to G>T on the opposing strand, as are the other cognate base pairs. (D) Established COSMIC signature MutationalPatterns analysis of scrambled control or *MT2A*-knockdown cells. (E) *De-novo* SBS signature analysis identified two novel signatures present in MT2A-KD cells, with standardized trinucleotide change frequencies plotted here. (F) Relative contribution of each de-novo MT2A-low SBS signature to MT2A-KD mutation patterns. (G) Fraction of mutations possibly attributable to *MT2A*-low SBS-ROS by MutationalPatterns analysis of TCGA PanCancer Atlas tumors. HGSC (OV) is highlighted in teal. (H) Fraction of mutations possibly attributable to *MT2A*-low SBS-BER by MutationalPatterns analysis of TCGA PanCancer Atlas tumors.

We hypothesize there may be an optimal knockdown level of *MT2A* for cancer cell proliferation and other oncogenic processes such as mutagenesis. Interestingly, while the MT2A-KD1 reduces *MT2A* mRNA more than MT2A-KD2, the mutational data suggested that there was a greater increase in the MT2A-KD2 condition. In the Dependency Map project portal DepMap, we noted that CRISPR-Cas9 mediated *MT2A* gene knockouts were strongly selective for slowed growth, whereas shRNA knockdown of *MT2A* formed a distribution of generally increased cell proliferation (**Supplementary Fig. S2A**). The top shared co-dependencies between the two screen types were other MT genes. This was validated by our observation of a significant decrease in proliferation with MT2A-KD1 (**Supplementary Fig. S2B**). These data are supportive that biological changes associated with MTs are dose dependent, and complete or stronger loss is deleterious to cell division. The MT2A-KD2 may exhibit more mutations than MT2A-KD1 due to more cell cycles during the 15 passages of the experiment. Our dose dependency hypothesis is consistent with the observation that tumors and cancer cell lines do not exhibit homozygous deletions of the MT gene cluster, but rather a heterozygous loss. Heterozygous losses in HGSC result in a 28% decrease in bulk mRNA expression on average (**Fig. 1C**), closer to our MT2A-KD2 model. There is a caveat that stresses likely present in clinical tumor RNA collection are inducers of MTs(50) and likely ameliorate true expression differences in clinically collected samples. Thus, while a complete loss of MTs may be detrimental for tumor development, heterozygous loss drives disease.

### Mutation signature analysis identifies oxidative mutations and impaired base excision repair in MT2A-knockdown cells

SMM-seq allows an assessment of potential mutational mechanisms as the mutations are randomly captured. Mutation spectra were quantified according to standard conventions of noting pyrimidine changes: cytosine to another nucleotide or thymine to another nucleotide. These SNVs were calculated for each genotype, with (**Supplementary Fig. S3A**) or without cadmium (**Fig. 2C**). Cadmium is reported to cause oxidative DNA damage in many studies(51–53). Oxidative DNA damage is often measured by 8-Oxo-2’-deoxyguanosine (8-oxo-dG) quantitation. 8-oxo-dG, if left unrepaired by base excision repair, yields a thymine transversion (G>T) upon DNA replication due to disrupted base pairing(54). With the standard single-base substitution (SBS) mutation convention(37), this would be calculated as a C>A change, the equivalent on the paired DNA strand of a G>T change. There was no increase in C>A changes in MT2A-KD cells passaged in cadmium chloride compared to vehicle control (**Supplementary Fig. S3A**, gold bars relative to blue bars), although a 26% increase in C>A was observed in control shScr cells passaged in cadmium chloride compared to vehicle control (\*\*\**P*<0.001, chi-squared test). The positive control with hydrogen peroxide induced mutations were also increased relative to vehicle control (**Fig. 2C, S3A** gray bar, \*\*\**P*<0.001, chi-squared test). However, vehicle MT2A-KD cells were C>A mutated at least three times the rate of those detected in vehicle shScr cells (**Fig. 2C** blue bars relative to black bar, \*\*\**P*<0.001, chi-squared test). Taken together, these data again indicate an epistatic relationship in which oxidative mutations caused by *MT2A* reduction are downstream of those caused by cadmium.

SBS signatures allow for an assessment of known and unknown mutation processes. The Catalogue Of Somatic Mutations In Cancer (COSMIC) database was utilized to score mutation signatures within the SMM-seq data and identify any *de novo* previously uncharacterized signatures (see Methods). Ovarian cancer, along with all other solid tumors, is characterized by SBS signature 1 and signature 5, which are clock-like signatures associated with aging. Ovarian cancer tumors, along with pancreatic and breast tumors, additionally harbor signature 3, a signature present in homology-directed repair deficient tumors. These signatures were not unique to any of the currently studied origin samples, as our analysis of spontaneous mutations filters out common mutations occurring in multiple samples and will remove signatures inherent to CAOV3 alone. Reactive oxygen species (ROS) provide acute, random, non-clonal mutagenesis, however, and would be predicted to be detected under the current experimental design. SBS signature 18 is largely attributed to ROS(37), and was indeed the most prevalent signature detected in this study (**Fig.2D**). However, overall, cadmium did not elevate SBS18 across all samples (**Supplementary Fig. S3B**), counter to the prevailing hypothesis that 8-oxo-dG is converted into spontaneous mutations after cadmium exposure. It is possible that our sublethal dose of cadmium is lower than other cadmium mutagenesis studies which may be above environmental exposure levels.

Interestingly, *de novo* trinucleotide signatures associated with MT2A-KD, or MT2A-low SBS, were detected in the nonnegative matrix factorization analysis (**Fig. 2E**). The first *de novo* signature involved largely C>A and C>T trinucleotide contexts (**Fig. 2E**). Cosine similarity analysis indicated these *de novo* signatures are most similar to SBS18 (cosine similarity = 0.82) (**Supplementary Fig. S4A**), a ROS-like signature, and SBS30 (cosine similarity = 0.76) (**Supplementary Fig. S4B**), a BER-defect signature. The MT2A-low SBS-BER signature was more prevalent than MT2A-low SBS-ROS within the SMM-seq MT2A-KD samples (**Fig. 2F**). Visually, MT2A-low SBS-ROS and MT2A-low SBS-BER appear to be a combination of both SBS18 and SBS30, along with a lower level of SBS-non-specific mutations, consistent with reduced HR(55). Assessing each *de novo* signature across TCGA-studied cancer types, substantial mutations were consistent with the MT2A-low SBS-ROS (**Fig. 2G**) and MT2A-low SBS-BER (**Fig. 2H**) signatures. OV, the HGSC study, was particularly high in the MT2A-low SBS-ROS signature (Fig. 2G). Parsing each TCGA cancer type into tumors with or without *MT2A* loss, five cancer types had significantly higher MT2A-low SBS-ROS signature in the *MT2A* loss samples (**Supplementary Fig. S5**). Signature analyses most clearly point to an involvement of ROS and failure of BER within *MT2A* loss tumor cells.

### Base excision repair rate is decreased in *MT2A*-knockdown cells

DNA damage alone is insufficient to yield mutagenesis; repair must be disrupted for a lesion to persist into replication and subsequent replacement with aberrant nucleotides. To analyze how the observed SMM-seq mutation spectrum resulted from decreased MTs, we undertook functional assays to assess BER. Unlike non-homologous end joining and homology-directed repair processes, BER does not allow for convenient reporter design in bacteria propagated DNA assays, such as plasmid reporter repair assays(56). For BER assessment, chemical lesions must be present on DNA and synthesized chemically for each repair assay evaluation. An assay designed to evaluate BER directly was recently innovated to allow for live cell testing by protecting the ends of the test molecule with mirror image _L_-DNA that enzymes cannot degrade, leaving the BER-repairable lesion intact long enough for BER-enzyme action in the nucleus(38,39). In the first step of BER, a lesion-specific DNA glycosylase, in this case uracil-DNA glycosylase (UNG) for the probe’s uracil incorporated DNA damage (**Fig. 3A**), flips the nucleobase out from the double helix structure. UNG releases the damaged base from DNA by cleaving the N-glycosidic bond, and leaves an apurinic/apyrimidinic (AP) site(57). Then the BER endonuclease APE1 cleaves the phosphodiester backbone, allowing the reporter CY5 fluorophore to detach, thereby activating fluorescence through distancing from the BER probe’s quenching agent, BHQ2 (**Fig. 3A**). By transfecting this probe into recipient cells with altered expression of *MT2A*, we were able to assess BER in live cells with varying levels of *MT2A*. The probe first travels to the cell’s endosomes and then a small portion escapes and travels to the nucleus (**Supplementary Movies S1,2**), wherein the nuclear CY5 signal was measured by microscopy after 24h of transfection (**Fig. 3B**). Counting cells with endocytosed probe yielded a result consistent with the SMM-seq mutation spectrum analysis: a decrease in repaired probe in the MT2A-KD cells (**Fig. 3C**). Again, there seemed to be a dose-dependent effect wherein the moderate knockdown clone MT2A-KD2 exhibited the more substantial decrease in BER signal, although both knockdowns were significant. MT2A-KD thus reduced BER efficiency.

**Figure 3.**
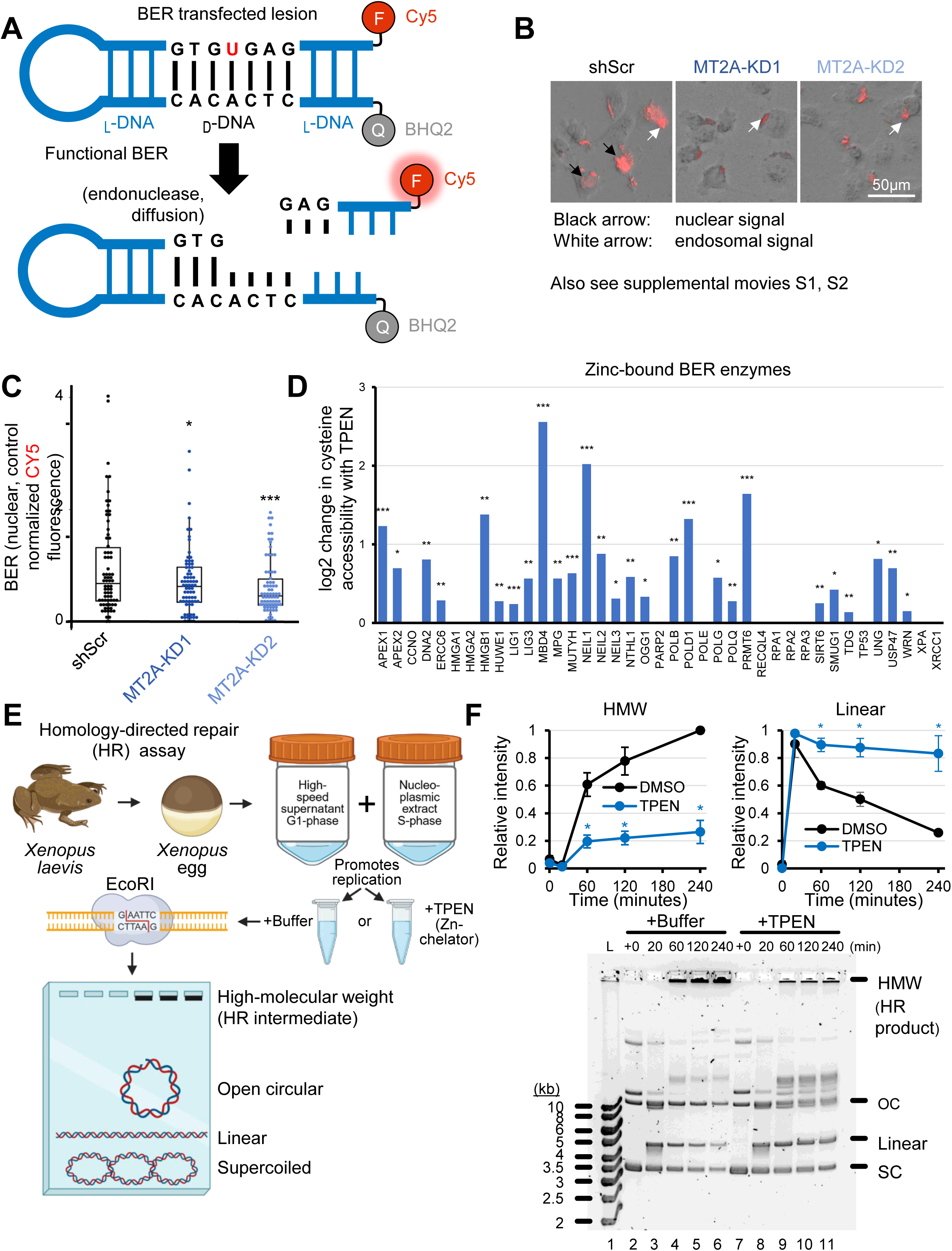
Defective base excision repair upon *MT2A* knockdown. (A) Depiction of the probe-based assay containing deoxyuracil-incorporated damaged DNA sequence, protected from cellular degradation by _L_-DNA flanking sequence. CY5 fluorescence is detectable upon strand cleavage by BER enzymes, separating the quenching moeity BHQ2 from the probe. (B) Live cell microscopy time course of probe fluorescence in CAOV3 cells with MT2A-KD or shScr control. (C) Quantitation of BER probe assay. \**P*<0.05, \*\**P*<0.01, \*\*\**P*<0.001 by Wilcoxon rank-sum test. (D) BER enzymes were queried in ZnCPT source data for changes in cysteine accessibility by TPEN addition, an indicator of zinc binding to queried protein. (E) Schematic of DNA repair assay to investigate HR using Xenopus egg extracts to repair DSBs generated by EcoRI and visualized via agarose gel electrophoresis. HMW represents high molecular weight branched DNA intermediates formed by HR. OC and SC are open circular and supercoiled plasmids, respectively. (F) Agarose gel and quantitation of *Xenopus* egg extract assessment of HR repair in the presence or absence of Zn chelator TPEN. \**P*<0.05 by t-test.

BER repairs oxidatively damaged DNA. Dozens of prior studies implicate oxidative stress as the primary mechanism of cadmium toxicity (51–53). Since MTs sequester cadmium, a straightforward hypothesis is that MTs protect against DNA damage by preventing cadmium’s mutagenic roles catalyzed by oxidative stress. Given that ROS and BER SBS signatures were hits for MT2A-KD, and that C>A [G>T opposing strand equivalent] spontaneous SNVs were a main mutagenic difference in MT2A-KD, we investigated oxidized DNA in *MT2A* deficient cells. OGG1, the glycosylase initiating BER during oxidative stress(58,59), is not significantly induced in protein expression by MT2A-KD nor cadmium stimulus (**Supplementary Fig. S6A**). While cadmium increased 8-oxo-dG in a dose dependent manner, 8-oxo-dG was not increased by MT2A-KD alone (**Supplementary Fig. S6B**). If oxidized guanine, 8-oxo-dG, is left unrepaired by BER, this results in a G>T mutation [equivalent to C>A](54). Given the observation that C>A SNVs are increased in MT2A-KD cells, but that this is not dependent on cadmium levels, the interpretation in the context of 8-oxo-dG data is that the cadmium-dependent increases in 8-oxo-dG are generally repaired well in the presence of adequate MT. However, in *MT2A* deficient cells, C>A mutations accumulate regardless of cadmium insult, and this is not further exacerbated by the presence of cadmium. Additionally, MT2A-KD cells treated with methyl methanesulfonate (MMS), a DNA alkylating agent, did not exhibit additional cell loss compared to scrambled control (**Supplementary Fig. S6C**). Together, this shows a lack of sensitivity to 8-oxo-dG and MMS with *MT2A* expression changes.

A direct molecular link from MTs to base excision repair is not immediately clear from the literature. A challenge to studying MTs is that, unlike kinases and ubiquitinases which covalently modify downstream proteins, MTs instead transfer labile metals: primarily zinc. Appreciation of possibly labile zinc binding sites across the proteome only came in 2025, with a careful mass spectrometry study using *in vitro* chelation of zinc by TPEN and protection chemistry (40). Approximately 10-15% of the proteome was found to bind zinc. The vast majority of BER enzymes contained Zn-bound cysteines and histidines, including OGG1 (**Fig. 3D**).

To investigate the effect of Zn on HR, we performed a homology-directed repair assay using *Xenopus laevis* egg extract. A combination of high-speed supernatant (HSS) and nucleoplasmic extract (NPE) was used to promote replication of a plasmid substrate containing a single EcoRI restriction site(60). Reactions were supplemented with Zn chelator TPEN or control buffer prior to the addition of EcoRI to generate a DNA double strand break with the restriction enzyme EcoRI. DNA intermediates were then isolated and resolved by agarose gel electrophoresis and analyzed for HR efficacy (**Fig. 3E**). We observed that HR, as measured by the formation of HR intermediates(60), was dramatically reduced with chelated Zn (**Fig.3F**). Thus, while BER defects in MT2A-KD were confirmed by mutation patterns and functional assays, other DNA repair pathways may also be disrupted by MT2A-KD and consequent Zn imbalance.

### *MT2A*-knockdown induces MYC and MAX target gene expression and cell stemness

Many of these Zn labile recipients of MT-donated zinc are zinc finger transcription factors, which require zinc to bind DNA. To assess how MTs affect transcriptional profiles, RNA-seq and EdgeR differential expression analysis was performed. 437 genes were upregulated and 355 genes were downregulated in MT2A-knockdown cells (**Fig. 4A**). The most downregulated transcription factor target groups were CCAAT/enhancer binding protein beta (CEBPB), a tumor suppressor which regulates G2/M cell cycle progression via kinase WEE1(61), and FOS, a transcription factor whose loss is associated with poorly differentiated ovarian cancers and worse prognosis(62) (**Fig. 4B**). Higher FOS can reduce the amount of circulating tumor cells in ovarian cancer mouse models(63), suggesting that FOS may suppress ovarian tumor cell dissemination and metastatic spread. The most upregulated pathway in MT2A-KD cells was MYC targets (**Fig. 4C**), consistent with our previous observation of increased MYC protein in MT2A-KD cells(11). The top co-expression transcriptional pathway regulator was REXO4 (**Fig. 4C**), an emerging cell cycle regulatory transcription factor regulating aggressive hepatocellular carcinomas(64).

**Figure 4.**
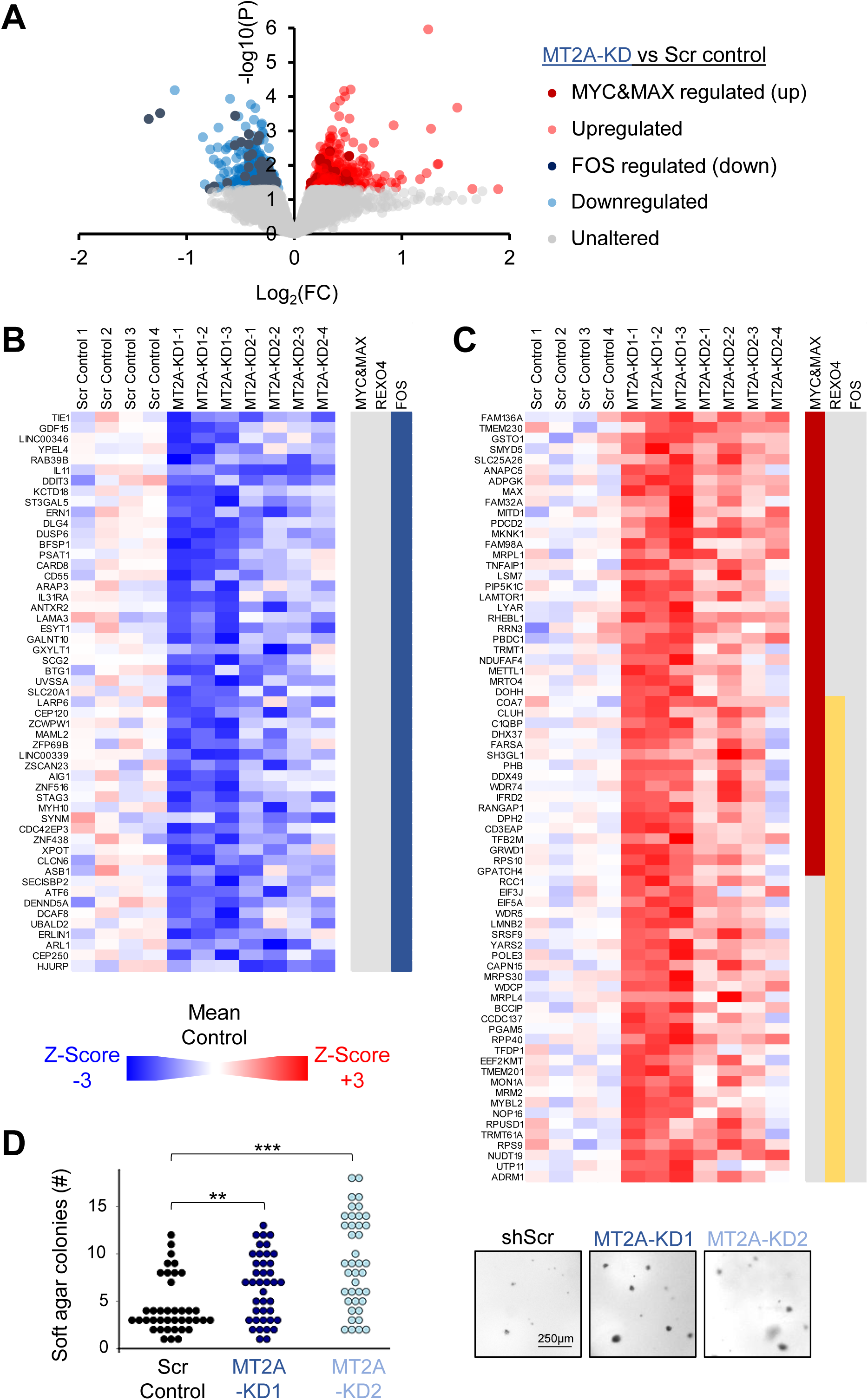
*MT2A* knockdown promotes a cell stemness program through upregulated MYC and MAX targets and downregulated FOS targets. (A) Volcano plot of transcribed genes affected by *MT2A*-knockdown, by bulk RNA-seq of CAOV3 cells. Transcription factors most enriched for dysregulated genes are indicated in panel legend. (B) FOS regulated transcripts that are downregulated (*P*<0.05) in *MT2A*-knockdown cells. (C) MYC and MAX regulated transcripts that are upregulated (*P*<0.05) in MT2A-knockdown cells. (D) Soft agar adhesion-free single-cell growth colony assay of *MT2A*-knockdown cells. \*\**P*<0.01, \*\*\**P*<0.001 by t-test.

To assess aspects of oncogenic transformation incurred by MYC/MAX functional enhancement in MT2A-knockdown cells, we performed a scratch wound assay, which examines if cancer epithelial cells acquire the ability to phenotypically transition to a mesenchymal state capable of faster migration. No differences were observed by scratch wound (**Supplementary Fig. S7A-B**). We tested the ability to form tumor spheres embedded in soft agar, a tumor initiating cell phenotype measured in anchorage independent growth conditions(65). MT2A-knockdown cells exhibited a marked increase in ability to form larger colonies in soft agar (**Fig. 4D**).

### *MT2A*-knockdown impairs mitochondrial homeostasis

To determine functional pathways beyond transcription factors dysregulated by MT2A-knockdown, we utilized shifted weighted annotation network (SWAN) analysis(32). MYC targets were again a highly impactful network, as was DNA replication. However, the most upregulated transcriptional network was mitochondrial translation (**Fig. 5A**). Since mitochondria regulate cell stemness and regulate reactive oxygen species related to the observed mutagenic profiles (Fig. 2), we opted to examine mitochondria in more detail in the context of MT2A-KD cells. Interestingly, mitochondrial morphology was more punctate and less contiguous, as assessed quantitatively as an increase in branching, for MT2A-knockdown cells (**Fig. 5B**). Functional impairment of mitochondria was then assessed. Mitochondrial membrane potential (MMP) was decreased in MT2A-KD, by a flow cytometric assay of TMRM fluorescence (**Fig. 5C**). Oxygen consumption rate, measured by Resipher and primarily an indicator of oxygen utilization within mitochondria, was decreased in MT2A-KD cells (**Fig. 5D**). General ionic destabilization may be implicated by the loss of zinc homeostasis in MT2A-KD cells. We observed hyperpolarization of the cell membrane by flow cytometry in MT2A-KD compared to shScr control (**Fig. 5E**). Taken together, the reduction of MT2A levels disrupts ion-dependent cell biological pathways, particularly mitochondria.

**Figure 5.**
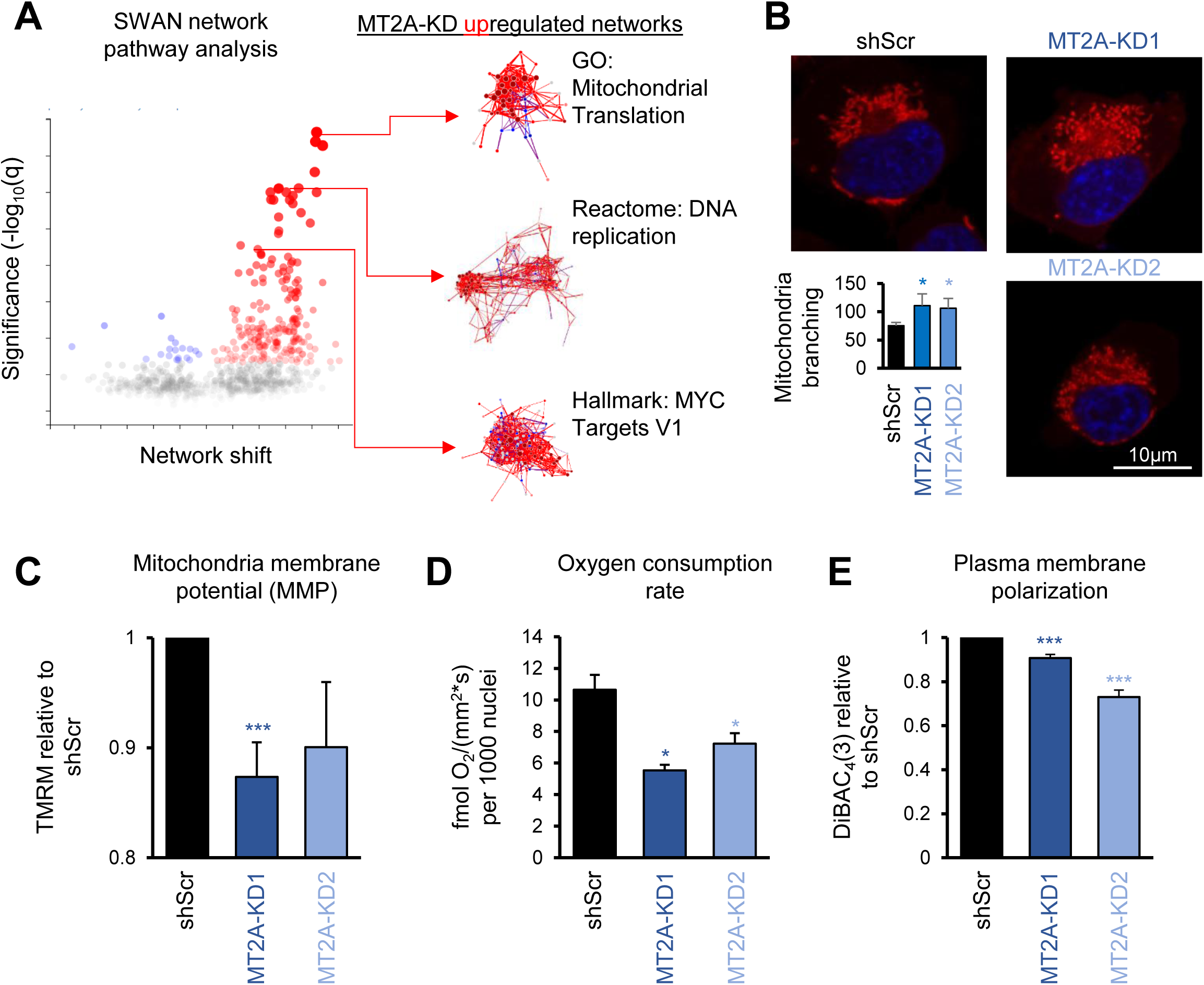
*MT2A* knockdown disrupts mitochondrial homeostasis. (A) SWAN network pathway analysis of transcriptome data of CAOV3 MT2A-KD cells compared to shScr control. Upregulated networks under molecular investigation are highlighted in network display panels. Red indicates upregulation, whereas blue indicates downregulation, with grey indicating non-significance. (B) CAOV3 transgenic cells were stained by MitoTracker Red, imaged by microscopy, and analyzed by the MitoTracker analyzer plugin in ImageJ. (C) Mitochondria membrane potential (MMP) of CAOV3 transgenic cells as assessed by TMRM flow cytometry. (D) Oxygen consumption rate of CAOV3 transgenic cells as assessed by Resipher assay. (E) Flow cytometry analysis of DiBAC_4_(3) stained CAOV3 transgenic cells assess to plasma membrane polarization. \**P*<0.05, \*\*\**P*<0.001 by t-test.

To confirm phenotype changes in an independent system, we utilized murine HGSC F318LOVi2 cells (see Methods for details). In this model, knockdown of *Mt2*, the murine ortholog of human *MT2A*, achieved approximately 60-70% reduction in mRNA and protein expression **(Fig. S8A-B)**. This corresponded to a functional decrease in ability to survive acute cadmium toxicity, as expected for MT suppression (**Fig. S8C**). MMP was decreased (**Fig. S8D**), similar in magnitude to the human cell case. Notably, γH2AX immunofluorescence shows an increase of punctae marking double-strand breaks in Mt2-KD, relative to shScr (**Fig. S8E**). Together, these data are consistent with our findings in the CAOV3 HGSC cell line and indicate coincident DNA damage and disrupted mitochondria function.

### Metabolomics indicate reduced pyruvate, amino acids, and imbalanced nucleobases in *MT2A*-knockdown

To better understand the connection between metabolic disruption and DNA repair deficiency in MT2A low cancer, we performed directed metabolism assays. By Seahorse, extracellular acidification rate (ECAR) was slowed in MT2A-KD1 cells following the addition of glucose and the mitochondrial ATP-synthesis inhibitor oligomycin (**Fig. 6A**). Polar liquid chromatography–mass spectrometry (LC-MS) metabolomics were performed on CAOV3 xenograft tumor samples. Consistent with the decrease in glycolysis as indicated by Seahorse assays, steady state pyruvate levels were decreased in *MT2A* knockdown tumors, despite no changes in glucose-6-phosphate (**Fig. 6B**). In accordance with an absence of acute peroxide sensitivity changes with *MT2A* knockdown (**Supplementary Fig. S9)**, no changes in ROS-scavenging glutathione were observed (**Fig. 6B**). Amino acids were largely reduced in levels in *MT2A* knockdowns, including asparagine, phenylalanine, tryptophan, isoleucine, serine, lysine, and tyrosine (**Fig. 6C**). Non-peptide and modified amino acids were also reduced, including most prominently N6-acetyl-L-lysine, which was the strongest reduced metabolite measured (**Fig. 6D**), in addition to N-acetyl-L-methionine and N-acetyl-L-alanine. N6-acetyl-L-lysine is the acetylated form of post-translationally modified lysine in proteins. Hypotaurine, but not taurine, was an exception in this pattern, and this oncometabolite was upregulated in MT2A-KD tumors (Fig. 6C).

**Figure 6.**
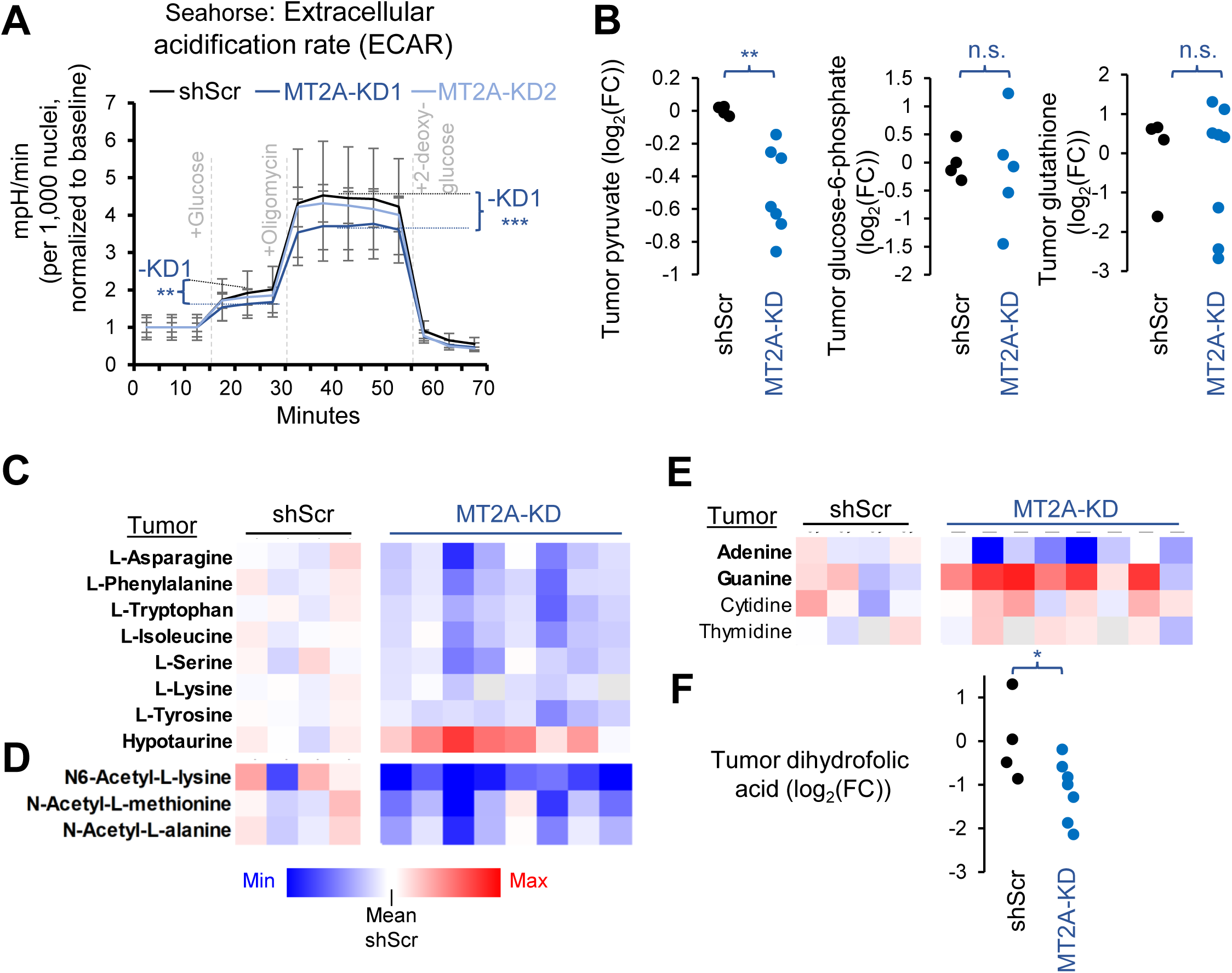
Metabolic rewiring in *MT2A* knockdown cells. (A) Seahorse extracellular acidification rate (ECAR) assay of MT2A-KD CAOV3 cells. (B-F) LC-MS metabolomics analysis of indicated substrates, from lysates acquired from transgenic CAOV3 intraperitoneal xenograft tumors. Bolded letters indicate statistical significance (*P*<0.05 by t-test). \**P*<0.05, \*\**P*<0.01, n.s. *P*>0.05.

Notably related to DNA metabolism, there was an imbalance of purine nucleobases at the steady state level in *MT2A* knockdown tumors compared to controls. Adenine was decreased, whereas guanine was increased (**Fig. 6E**). Dihydrofolic acid, a necessary precursor for purine and pyrimidine synthesis in the folate metabolic pathway, was the second most strongly downregulated metabolite in *MT2A* knockdown tumors (**Fig. 6F**). Folate metabolism notably interacts with mitochondria metabolism to allow for the synthesis of adenine and guanine(66). With a properly regulated cell cycle, cells will slow transitions during S-phase due to nucleobase imbalance(67). However, in these p53-mutant, MYC-driven ovarian cancer cells, cells may proceed through the cell cycle in the absence of correctly incorporated nucleotides. This hypothesis is consistent with the increased mutations observed with passaging *MT2A*-deficient cells (Fig. 2).

## Discussion

Here, we provide evidence of a novel central regulator of human cell mutagenesis: metallothioneins (MTs). We show that loss of *MT2A*, a frequently deleted MT gene in HGSC(11), drives interconnected phenotypes including increased mutagenesis, decreased DNA repair, metabolic dysfunction, and enhanced cell stemness. Using controlled *MT2A* knockdown models in HGSC cells, we demonstrate that *MT2A* loss is sufficient to increase oxidative DNA mutations, independent of cadmium exposure, underscoring a previously unappreciated role for *MT2A* in maintaining genomic integrity. Our application of single-molecule mutation sequencing revealed that *MT2A*-deficient cells accumulate fixed somatic mutations with mutation signatures most closely aligning with BER defects and oxidative stress. Functional BER assays in live cells indicated a deficiency in BER with reduced *MT2A* levels. These results establish a direct link between *MT2A* and genome stability in human cells.

*MT2A* loss disrupts mitochondrial morphology, function, and metabolism. We observed increased mitochondrial fragmentation, decreased membrane potential, and reduced oxygen consumption rates in *MT2A*-deficient cells. Metabolomics analyses revealed a broad decrease in amino acid pools, decreased pyruvate, and altered purine metabolism, collectively indicating impaired anabolic metabolism and overall metabolic slowing. Notably, acetylated and non-acetylated amino acids were decreased whereas hypotaurine levels were elevated, consistent with impaired mitochondrial and cellular regulation. Hypotaurine stabilizes HIF-1 and acts as an oncometabolite in glioma(68) and colorectal cancers(69). These mitochondrial and metabolic disruptions may further limit the energetic and cofactor resources required for DNA repair, creating a permissive environment for mutagenesis(70). Moreover, these results suggest that *MT2A*-deficient tumors may harbor unique metabolic vulnerabilities, which could be therapeutically exploited(71).

*MT2A* deficiency was found to promote stemness programs. *MT2A* knockdown cells exhibited upregulation of MYC and MAX targets and downregulation of FOS targets, coupled with enhanced single-cell colony formation in adhesion-free soft agar(65). *FOS*, which in some cancers is a proto-oncogene, acts differently in ovarian cancer (**Fig. 4B**). Higher FOS reduces metastatic cell populations in HGSC(63), and lower FOS is poorly prognostic in HGSC(62). This observation suggests that genomic instability and metabolic dysregulation may feed into transcriptional programs that favor a stem-like, tumorigenic state. We propose that mitochondrial dysfunction, decreased DNA repair, and metabolic reprogramming act together to reinforce stemness in *MT2A*-deficient cells, which is linked to more aggressive cancer phenotypes and poor outcomes in HGSC.

MT2A most likely sustains DNA repair capacity through its role in zinc homeostasis. Zinc is essential for the structural and catalytic activity of numerous DNA repair enzymes, particularly those in the base excision repair pathway (**Fig. 3D**)(19,72). Our analyses indicate that *MT2A*-deficient cells exhibit impaired repair despite unaltered OGG1 expression and 8-oxo-dG levels, suggesting that the functional activity of BER enzymes is compromised, likely due to alterations in zinc homeostasis. This aligns with prior studies demonstrating that perturbations in cellular zinc homeostasis impair BER and increase mutation accumulation(72). These findings are also consistent with an *in vivo* zinc deficient diet study showing an increase in 8-oxo-dG and repair protein expression in rats(73), amongst other studies (reviewed by Sharif *et al*(74)), suggesting that disrupting Zn homeostasis leads to aberrant DNA repair. Together, these data support a model in which MT2A helps maintain genomic integrity not by directly preventing oxidative damage, but by sustaining zinc-dependent DNA repair processes. While new technologies to study Zn *in vivo* are required, further experiments to directly test zinc transfer and disruption by MT2A will be of interest in future studies.

Collectively, our findings establish a mechanistic framework linking *MT2A* loss to multiple hallmarks of cancer: genomic instability, metabolic reprogramming, mitochondrial dysfunction, and stemness. The ability of MT2A to coordinate DNA repair and maintain mitochondrial health highlights its central role in preserving genomic stability in HGSC.

## Conclusions

In summary, *MT2A* loss is identified for the first time as a key driver of genomic instability. Mutational patterns with *MT2A* suppression were consistent with ROS-like and BER-defective signatures. *MT2A* loss further led to metabolic dysfunction and stem-like phenotypes in HGSC. These findings integrate DNA repair, mitochondrial biology, and metabolism, revealing a previously unappreciated mutagenic factor in cancer.

## Supplemental Data

**Supplementary Figure S1. Mutational analysis in the context of passaging in CdCl_2_.** (A) Schematic of genomic DNA sample collection for use in SMM-seq. (B) Quantitation of any SNV from SMM-seq data with indicated statistics derived from Fisher’s exact tests.

**Supplementary Figure S2. Dose-sensitive growth changes with MT2A-KD.** (A) DepMap output of cancer cell line screens from RNAi screens and CRISPR-Cas9 knockout screens, with a null effect of a gene knockdown centered at 0. Positive numbers indicate faster growth whereas negative numbers indicate impaired growth and cancer dependency. (B) Proliferation assay of CAOV3 MT2A-KD cells compared to shScr.

**Supplementary Figure S3. Spontaneous single nucleotide variant analysis in the context of CdCl_2_**. (A) Specific nucleotide changes SNVs per Mb of DNA analyzed by SMM-seq, including cadmium treatment. (B) SBS signatures as a fraction of total mutations, ranked by those most present in vehicle samples and compared to CdCl_2_ treated (excluding peroxide) samples.

**Supplementary Figure S4. Established SBS signatures cosine most-similar to *MT2A*-low SBS signatures.** (A) Cosine similarity analysis of SBS MT2A-low-ROS indicated the highest similarity to SBS18, plotted in trinucleotide context here. (B) Cosine similarity analysis of SBS MT2A-low-BER indicated the highest similarity to SBS30, plotted in trinucleotide context here.

**Supplementary Figure S5. Novel signature attributable mutations per cancer type by *MT2A* loss.** MutationalPatterns analysis of TCGA tumors for the MT2A-low (ROS-like) *de-novo* signature, indicating the percent of SNV mutations that may be attributable to the *de-novo* signature. Colored box plots indicate heterozygous *MT2A* loss, and gray represents no *MT2A* loss.

**Supplementary Figure S6. No 8-oxo-dG or MMS sensitivity changes with MT2A-KD.** (A) Western blots from samples with indicated genetics and antibodies (mean +/- s.e.m. from 2 experiments). (B) Quantitation of an ELISA-based 8-oxo-dG assay (mean +/- s.e.m. from 4 experiments), normalized to shScr 0µM CdCl_2_. *P<0.05; **P<0.01 by t. test compared to 0µM CdCl_2_. (C) Cell loss assay by nuclei count (Hoechst 33342) with increasing doses of methyl methanesulfonate (MMS) DNA alkylating agent treatment. n.s. indicates *P*>0.05 by t-test.

**Supplementary Figure S7. Scratch wound assay in MT2A-KD cells.** (A) Quantitation of scratch assay. P>0.05 by t-test. (B) Representative images of the scratch assay.

**Supplementary Figure S8. F318LOVi2 Mt2-KD metabolic changes.** All panels in this figure use the F318LOVi2 murine cell line. (A) RT-qPCR confirmation of *Mt2-*KD. (B) Immunofluorescence localization of Mt2 in control or knockdown cells. (C) Nuclei count assay following 72-hr CdCl_2_ treatment to verify a functional Mt2 knockdown. (D) Flow cytometry analysis of mitochondria membrane potential by TMRM. (E) Immunofluorescent staining of γH2AX associated DNA damage and Hoescht 33342. Cellpose-mediated segmentation for punctae quantitation is shown relative to shScr normalization per experiment. *P<0.05, **P<0.01, ***P<0.001by t-test.

**Supplementary Figure S9. No peroxide sensitivity changes in MT2A-KD.** Proliferation assay by nuclei count following 72-hour peroxide treatment in a dose-dependent manner. n.s. indicates *P*>0.05 by t-test.

**Supplementary Table S1.** Primer sequences used

**Supplementary Table S2.** Excel workbook of raw VCFs of spontaneous SNV calls per sample from SMM-seq

**Supplementary Table S3.** Spontaneous SNV summary from SMM-seq

**Supplementary Table S4.** *De novo* SBS signatures of MT2A-low from SMM-seq

**Supplementary Movie S1.** Hourly imaged movie of shScr control CAOV3 cells transfected with live cell BER assessment reagent.

**Supplementary Movie S2.** Hourly imaged movie of MT2A-KD2 CAOV3 cells transfected with live cell BER assessment reagent.

## Acknowledgments

We thank all private donors to oncology research, particularly Matt Prisby for organizing the Sheryl Prisby gynecologic oncology fund, for their individually perhaps small yet cumulatively enormous contributions to advancing cancer research and health. We thank Hyland Gonzalez and Kristy Thomas for their guidance on the Resipher assay. We thank Li Li for her help with confocal imaging and Jacob Kendrick and flow core members for their help with flow cytometry. We thank all Delaney lab members and collaborators who reviewed the initial and corrected manuscripts.

## Author contributions

MMA (Conceptualization, Investigation, Data curation, Methodology, Software, Funding acquisition, Formal Analysis), ACR (Conceptualization, Investigation, Data curation, Methodology, Formal Analysis), DE (Conceptualization, Investigation, Data curation, Methodology, Formal Analysis), EV (Conceptualization, Investigation, Data curation, Methodology, Formal Analysis), GF (Investigation, Methodology), HRG (Conceptualization, Investigation, Data curation, Methodology, Formal Analysis), MAC (Conceptualization, Investigation, Data curation, Methodology, Formal Analysis), WYZ (Conceptualization, Investigation), IK (Investigation, Methodology), IB (Investigation, Methodology), SO (Investigation, Methodology), RH (Investigation, Methodology), DD (Investigation, Methodology), Cd (Investigation, Methodology, Formal Analysis), AYM (Investigation, Methodology), CB (Investigation, Methodology), SS (Investigation, Methodology), REPA (Conceptualization, Investigation, Data curation, Methodology), YKP (Methodology), JZ (Methodology), ZY (Methodology), TCR (Methodology, Investigation, Formal Analysis), DMT (Methodology), SG (Conceptualization, Methodology, Supervision), BO (Conceptualization, Supervision), DJ (Conceptualization, Investigation, Data curation, Methodology, Software, Funding acquisition, Formal Analysis), JHH (Conceptualization, Methodology, Supervision), DTL (Conceptualization, Data curation, Methodology, Funding acquisition, Formal Analysis, Supervision), JTS (Conceptualization, Investigation, Data curation, Methodology, Funding acquisition, Formal Analysis, Supervision), JRD (Conceptualization, Investigation, Data curation, Methodology, Software, Funding acquisition, Formal Analysis, Resources, Supervision). All authors contributed to the writing and approval of the final manuscript.

## Data Availability

Raw data for SMM-seq and spectrum analysis have been deposited in SRA under Bioproject PRJNA1213734. Transcriptomic analysis raw sequencing data are available within SRA under Bioproject PRJNA1428857 and the raw count data is deposited in GEO under series GSE322490. Code associated with this study is available at https://github.com/jrdelaney/Published_Code/.

