## Supplementary Figures1-9 for "Metallothionein loss in cancer cells contributes to increased mutations through defective DNA repair and metabolic imbalance"

### Supplementary Fig. S1: Mutational analysis in context of passaging in $CdCl_2$

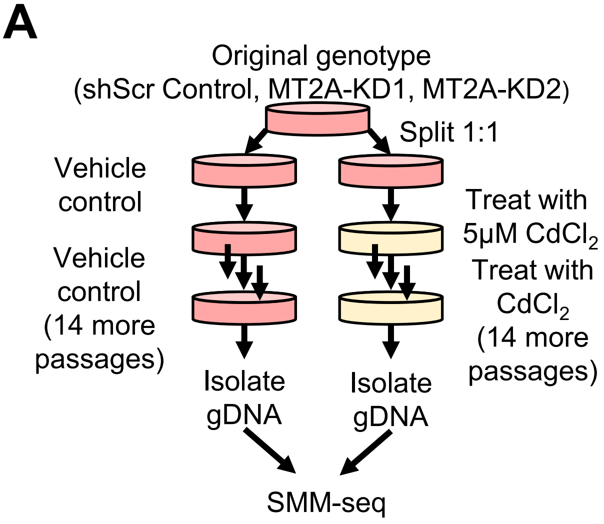


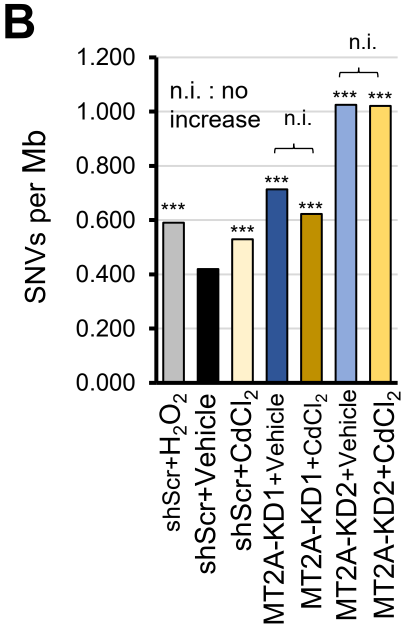

**Supplementary Fig. S2: Dose-sensitive growth changes with MT2A-KD**

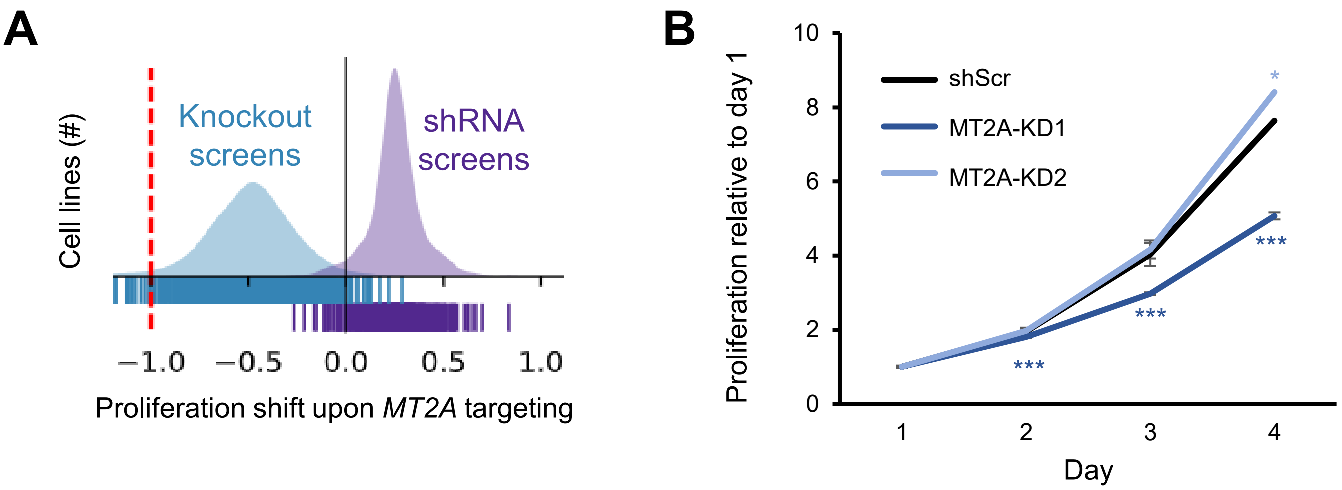

**Supplementary Fig. S3: Spontaneous single nucleotide variant analysis in the context of $CdCl_2$**

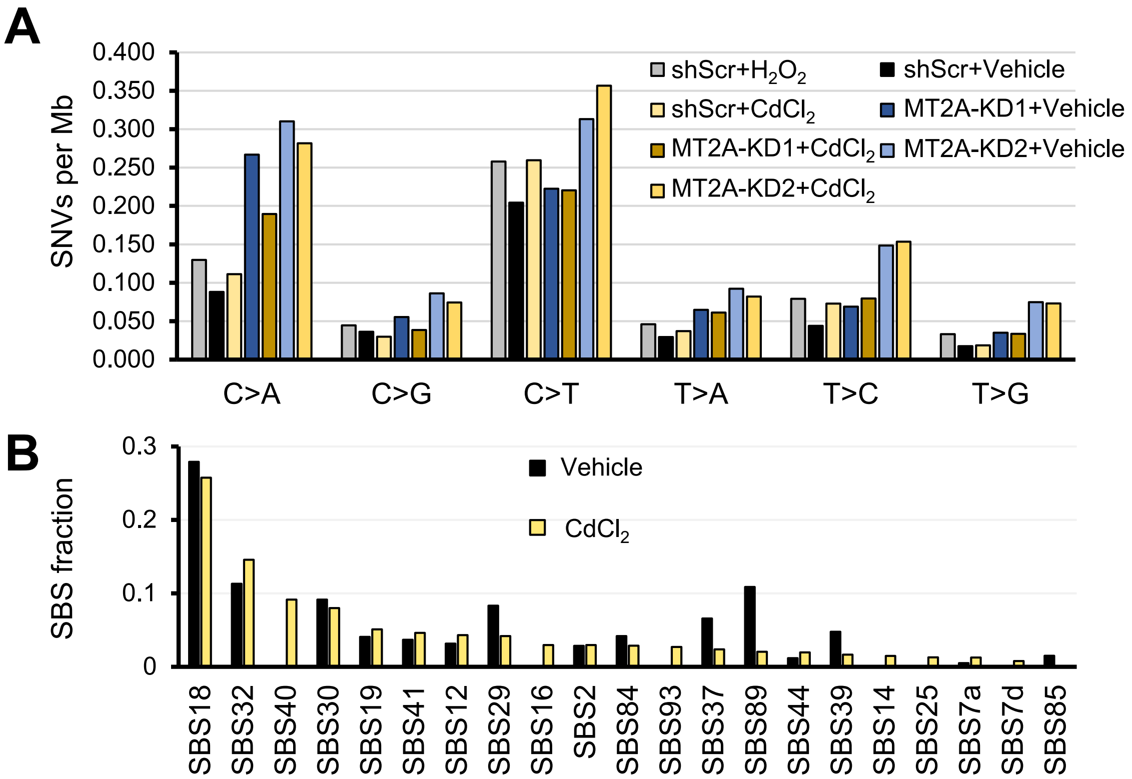

#### Supplementary Fig. S4:
#### Established SBS signatures cosine most-similar to *MT2A*-low SBS signatures

**A** MT2Alow SBS-ROS:SBS18 (Possibly damage by reactive oxygen species)
cosine similarity = 81.8%

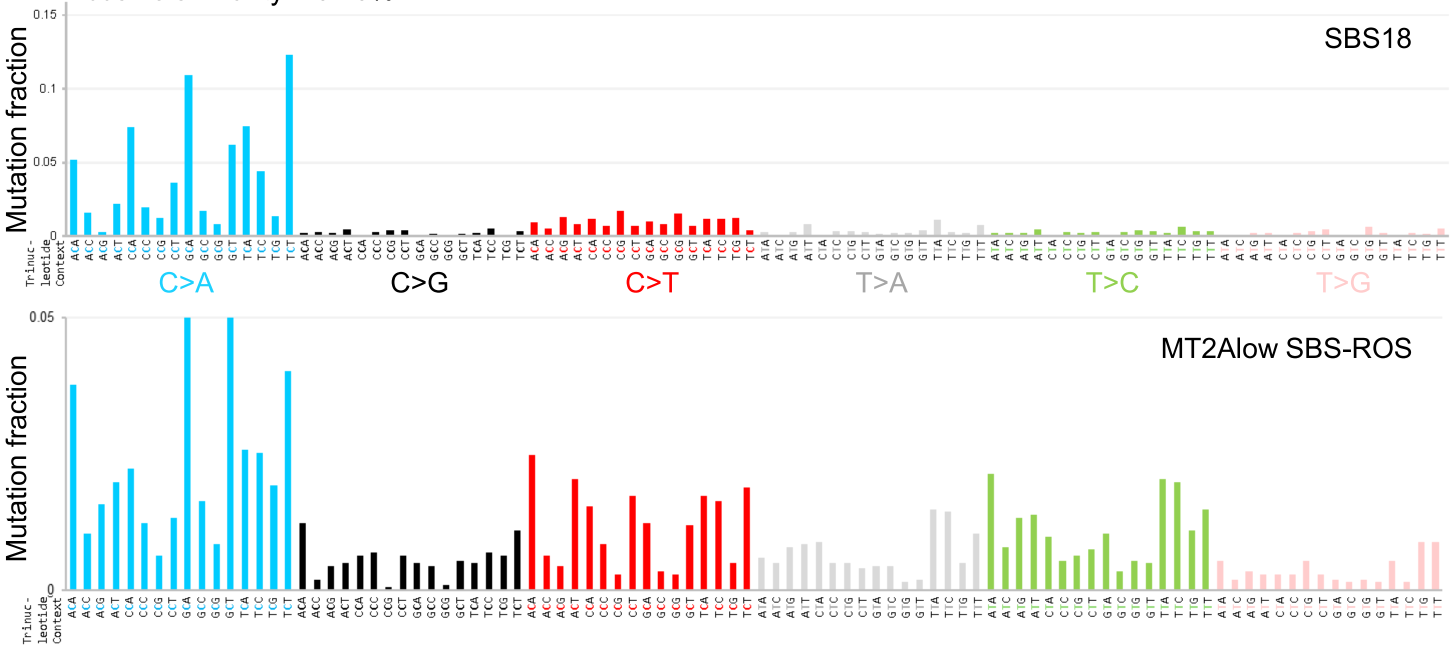


**B** MT2Alow SBS-BER:SBS30 (base excision repair due to inactivating mutations in *NTHL1*)
cosine similarity = 75.6%

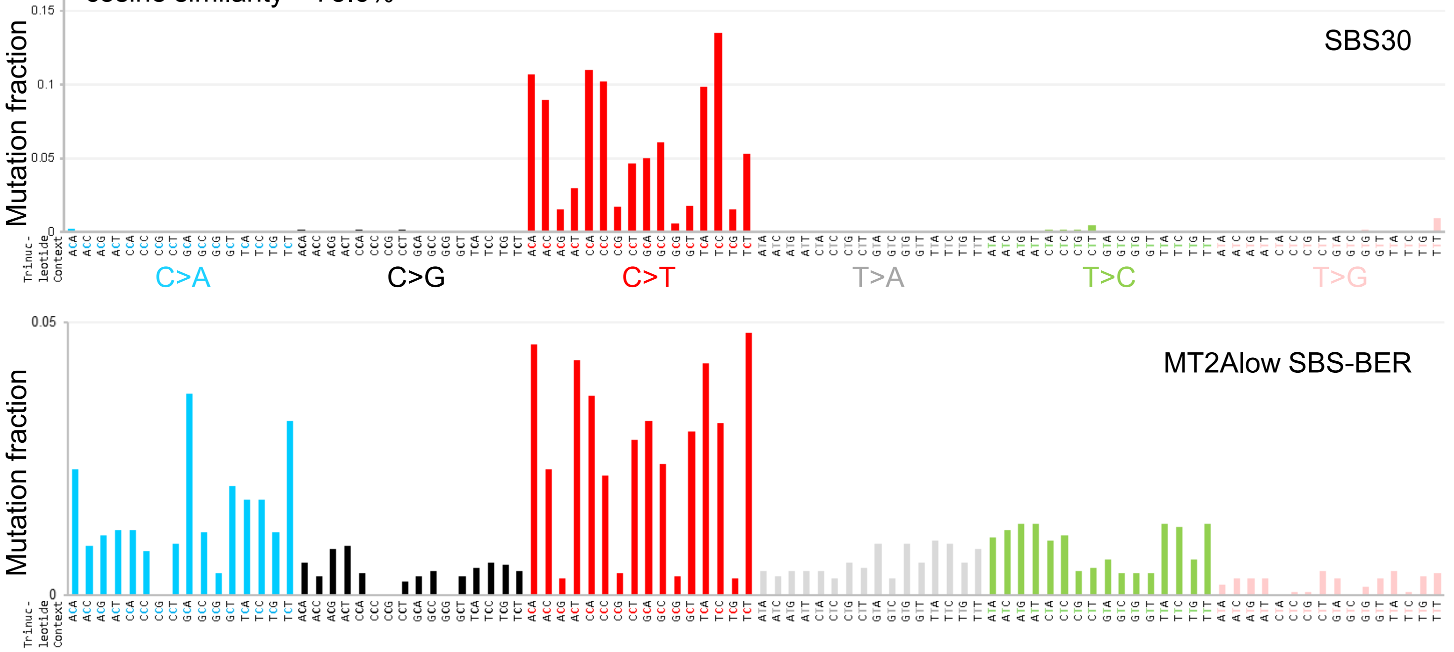

**Supplementary Fig. S5:**
**Novel signature attributable mutations per cancer type by *MT2A* loss**

MT2A-low *de-novo* (ROS-like) SBS signature MutationPatterns analysis of TCGA tumors
Colored: *MT2A* heterozygous loss
Gray: No *MT2A* loss

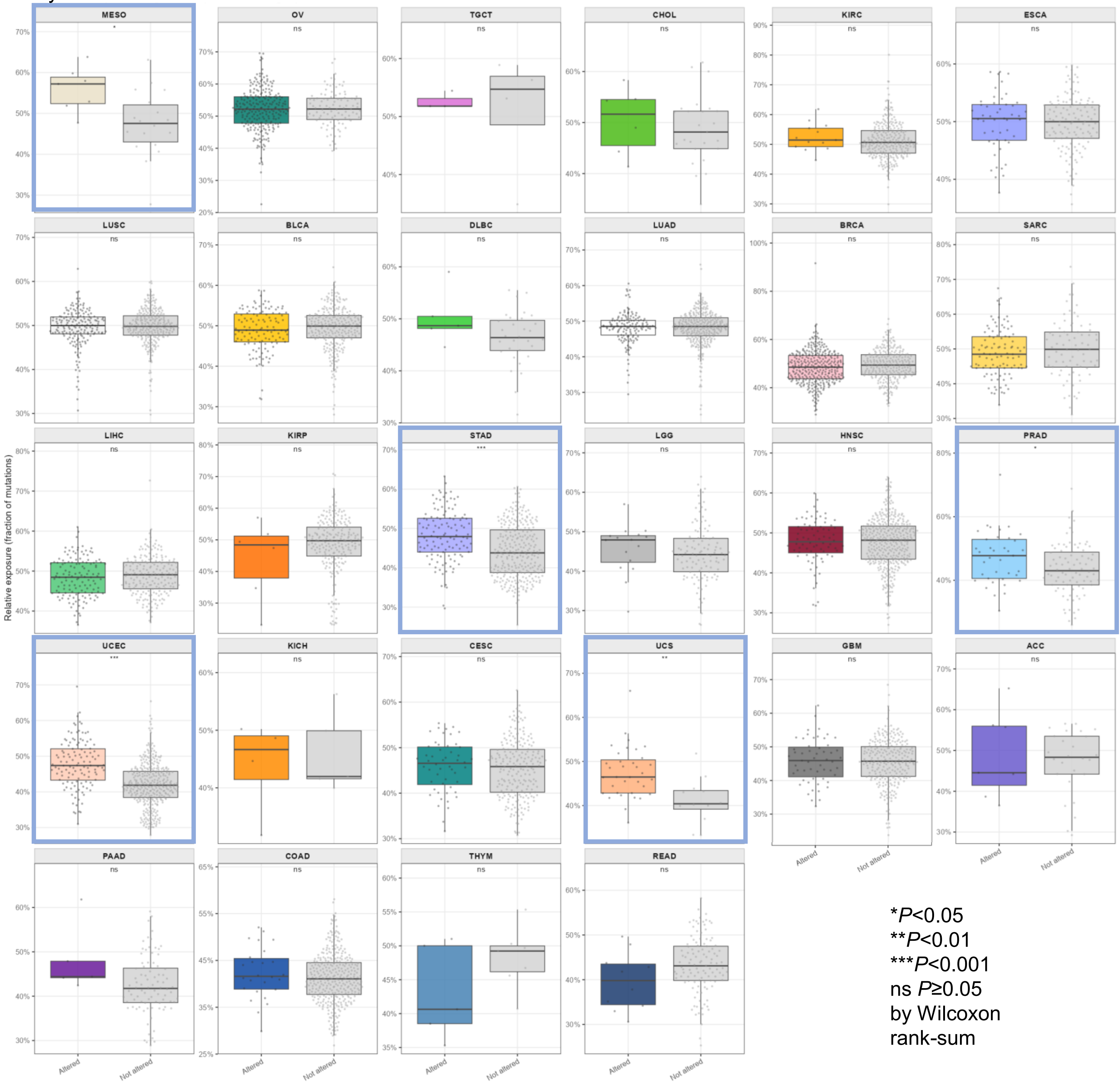


*$P<0.05$
**$P<0.01$
***$P<0.001$
ns $P\geq0.05$
by Wilcoxon rank-sum

**Supplementary Fig. S6: No 8-oxo-dG or MMS sensitivity changes with MT2A-KD**

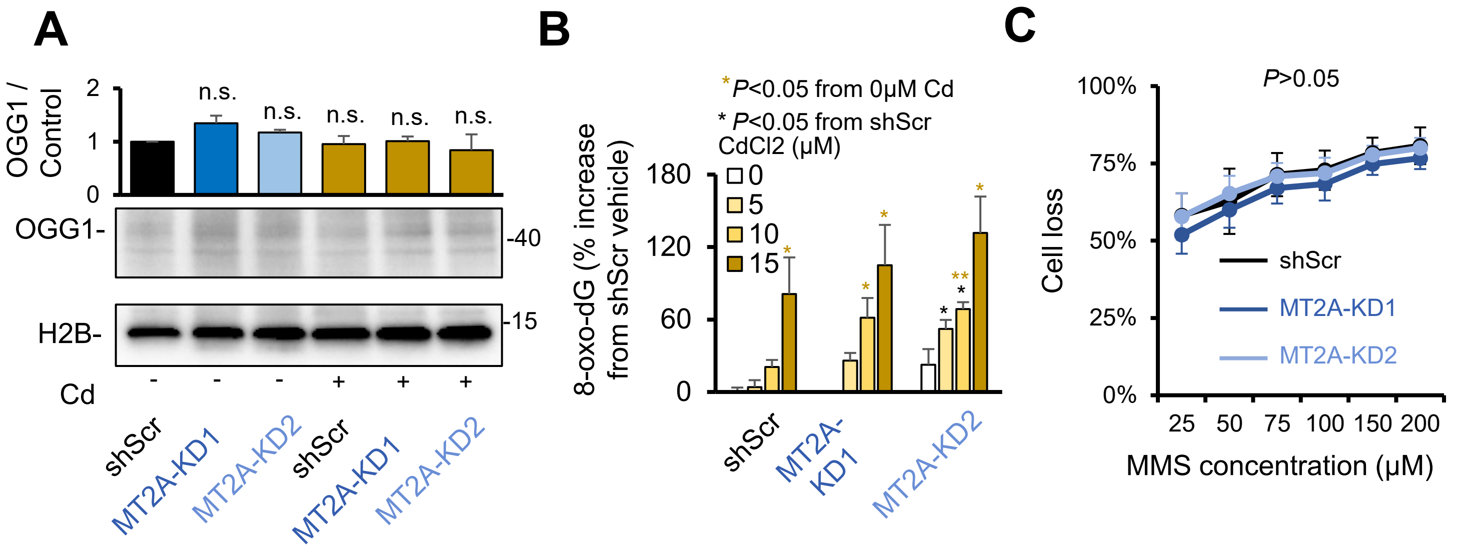

**Supplementary Fig. S7: Scratch wound assay in MT2A-KD cells**

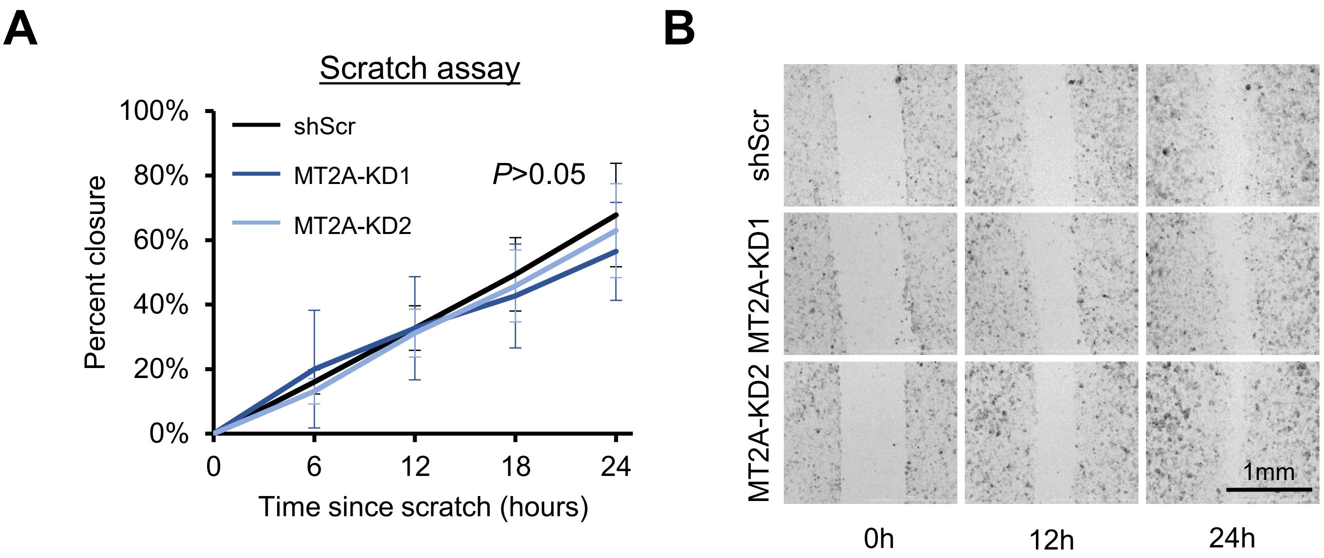

**Supplementary Fig. S8: F318LOVi2 Mt2-KD metabolic changes**

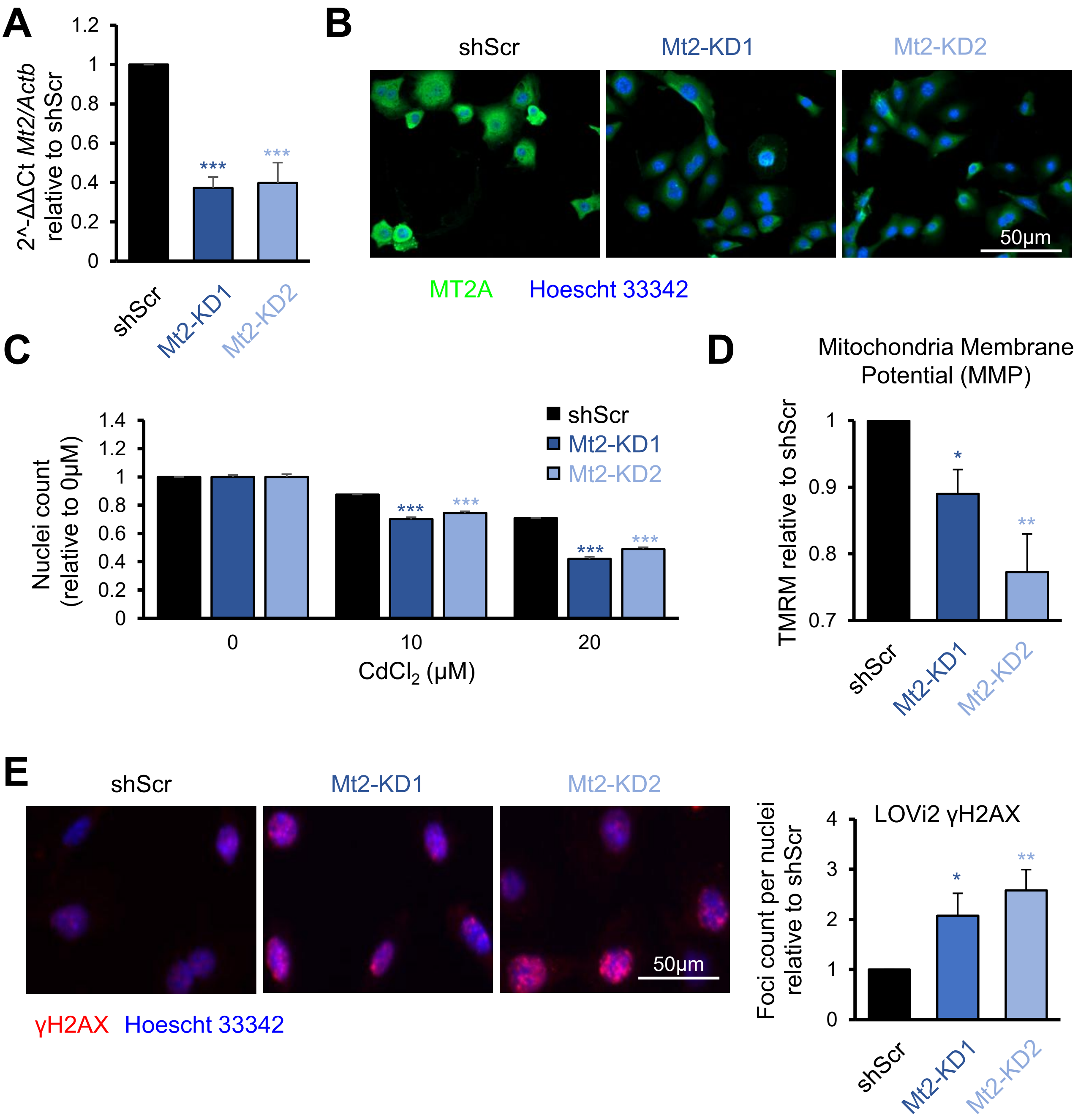

**Supplementary Fig. S9: No peroxide sensitivity changes in MT2A-KD**

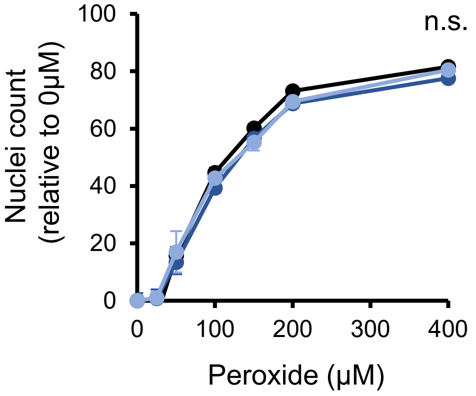
